# Lactate Accelerates Mouse ES Cell Differentiation Towards the XEN Lineage

**DOI:** 10.1101/2020.12.14.422783

**Authors:** Mohamed I. Gatie, Tyler T. Cooper, Gilles A. Lajoie, Gregory M. Kelly

## Abstract

Metabolism plays a crucial role for cell survival and function; however, recent evidence has implicated it in regulating embryonic development. The inner cell mass undergoes orchestrated cellular divisions resulting in the formation of embryonic stem cells and extraembryonic endoderm (XEN) cells. Concomitantly, changes in the metabolic profile occurs during development and are well-documented in the embryonic lineages. However, a comprehensive multi-omics analysis of these features in XEN cells remains lacking. We observed that feeder-free XEN cells exhibited high sensitivity to glycolytic inhibition in addition to maintaining elevated intra- and extracellular lactate levels. XEN cells maintain high lactate levels by increased LDHA activity and re-routing pyruvate away from the mitochondria. Importantly, exogenous lactate supplementation or promoting intracellular lactate accumulation enhances XEN differentiation *in vitro*. Our results highlight how lactate contributes to XEN differentiation in the mammalian embryo and may serve to enhance reprogramming efficiency of cells used for regenerative medicine.

**Highlights:**

- Feeder-free XEN cells exhibit high sensitivity to glycolytic inhibition
- Distinct transcriptomic, proteomic and metabolomic profile exists between feeder-free ES and XEN cells
- Elevated intracellular and extracellular lactate is observed in feeder-free XEN cells
- Lactate enhances feeder-free XEN differentiation *in vitro*

## Introduction

The early mammalian blastocyst is comprised of two distinct layers: the inner cell mass (ICM) and trophectoderm. The ICM houses embryonic stem (ES) cells and primitive endoderm cells, and the latter differentiate into parietal and visceral endoderm cells. Collectively primitive, parietal and visceral endoderm cells make up the extraembryonic endoderm (XEN) lineage. Various ES cells have been cultured at different developmental time-points (Evans and Kaufman, 1981), each requiring specific conditions. For instance, preimplantation ICM cells represent a “naïve” state and can be artificially maintained under LIF and serum/BMP (Nichols et al., 1990) or defined conditions of LIF-2i in N2B27 media where dual inhibition of glycogen synthase kinase 3 and mitogen-activated protein kinase pathways maintain pluripotency (Ying et al., 2008). However, the post-implantation or “primed” epiblast stem cells, which are poised to give rise to the embryo proper, require Activin A and FGF2 in culture (Brons et al., 2007). In addition to differences in culture conditions, both naïve and primed states differ in their expression profile of pluripotency core genes, DNA methylation, histone modification status, clonogenicity and chimera contribution (Davidson et al., 2015). XEN cells, which contribute to both embryonic and extraembryonic endoderm tissue (Nowotschin et al., 2019), have been derived by isolation from blastocyst (Niakan et al., 2013), differentiated using exogenous factors (Anderson et al., 2017; Cho et al., 2012; Soprano et al., 2007), reprogrammed from fibroblast cells (He et al., 2020; Parenti et al., 2016), or form after the overexpression of *Gata4/6* (Fujikura et al., 2002; Shimosato et al., 2007) or *Sox17* (McDonald et al., 2014). In addition, evidence indicates that the metabolic state between embryonic and extraembryonic lineages differs, suggesting that metabolism may be sufficient to influence lineage commitment.

Naïve ES cells exhibit a bivalent metabolic profile, relying on glycolysis and oxidative phosphorylation (OXPHOS) to generate energy, while epiblast stem cells are exclusively glycolytic, despite displaying a mature mitochondrial ultrastructure (Zhou et al., 2012). Although the majority of studies have focused on naïve and primed ES cells, recent reports on extraembryonic cells show they are reliant on OXPHOS metabolism (Choi et al., 2020), with a similar mitochondrial ultrastructure to epiblast stem cells (Seo et al., 2020; Zhou et al., 2012). However, details on the comprehensive intracellular profile of metabolites and their role during differentiation remains unknown.

Small metabolites have garnered attention with their role in maintaining pluripotency, promoting differentiation and enhancing reprogramming efficiency of induced pluripotent stem cells (Mathieu and Ruohola-Baker, 2017; Tsogtbaatar et al., 2020; Zhang et al., 2012). These small compounds play integral roles in energy production and epigenetic modifications. For instance, short-chain and saturated fatty acids can modulate histone deacetylases and the epigenetic landscape and thus are implicated in enhancing the efficiency of stem cell differentiation (Mali et al., 2010; Yanes et al., 2010). Also, the depletion of threonine or loss of threonine dehydrogenase (TDH) impacts stemness (Wang et al., 2009), while supplementation of L-threonine maintains pluripotency (Ryu and Han, 2011). Therefore, naturally occurring metabolites play an important role in regulating pluripotency and differentiation of stem cells by modulating the metabolic and epigenetic landscape.

In this report we used a multi-omics approach to identify metabolic changes in cells of the ICM and have further characterized the metabolic pathways involved in regulating XEN lineage commitment. Overall, we observed a significant reduction in the survival of feeder-free XEN cells following glycolytic inhibition. Transcriptomic and proteomic profiling implicate lactate metabolism in regulating the differentiation towards the XEN lineage. Strikingly, lactate was enriched intra- and extracellularly in feeder-free XEN cells, and promoting intracellular lactate accumulation or supplementing with exogenous L-lactate enhanced XEN induction. Together our results elucidate an unacknowledged role for lactate in cell differentiation and fate commitment in the early mammalian embryo.

## Results

### ES cells are less sensitive to glycolytic inhibition than XEN cells

A recent report characterized the metabolic profile of XEN cells as reliant on OXPHOS metabolism (Choi et al., 2020). Albeit informative, this study derived and maintained XEN cells on inactivated mouse embryonic fibroblasts, which are oxidative in nature (Folmes et al., 2011) and are known to reduce the reliance of pluripotent stem cells on glycolysis (Gu et al., 2016). To address if culturing methods influence the metabolic profiles, ES-E14TG2a embryonic stem cells (ES cells) and extraembryonic endoderm cells (XEN cells) (Kunath et al., 2005) were cultured under feeder-free conditions in media containing 50mM 2-Deoxy-D-glucose (2-DG; glycolysis inhibitor) or 2.5μM oligomycin (OXPHOS inhibitor), and results were compared with cells grown under control conditions (Figure 1). ES cells displayed domed-homogenous colonies while XEN cells were single-celled, with two distinct morphologies present in culture, spindle-like or round and refractile in nature (Figure 1A, Figure S1A).

**Figure 1.**
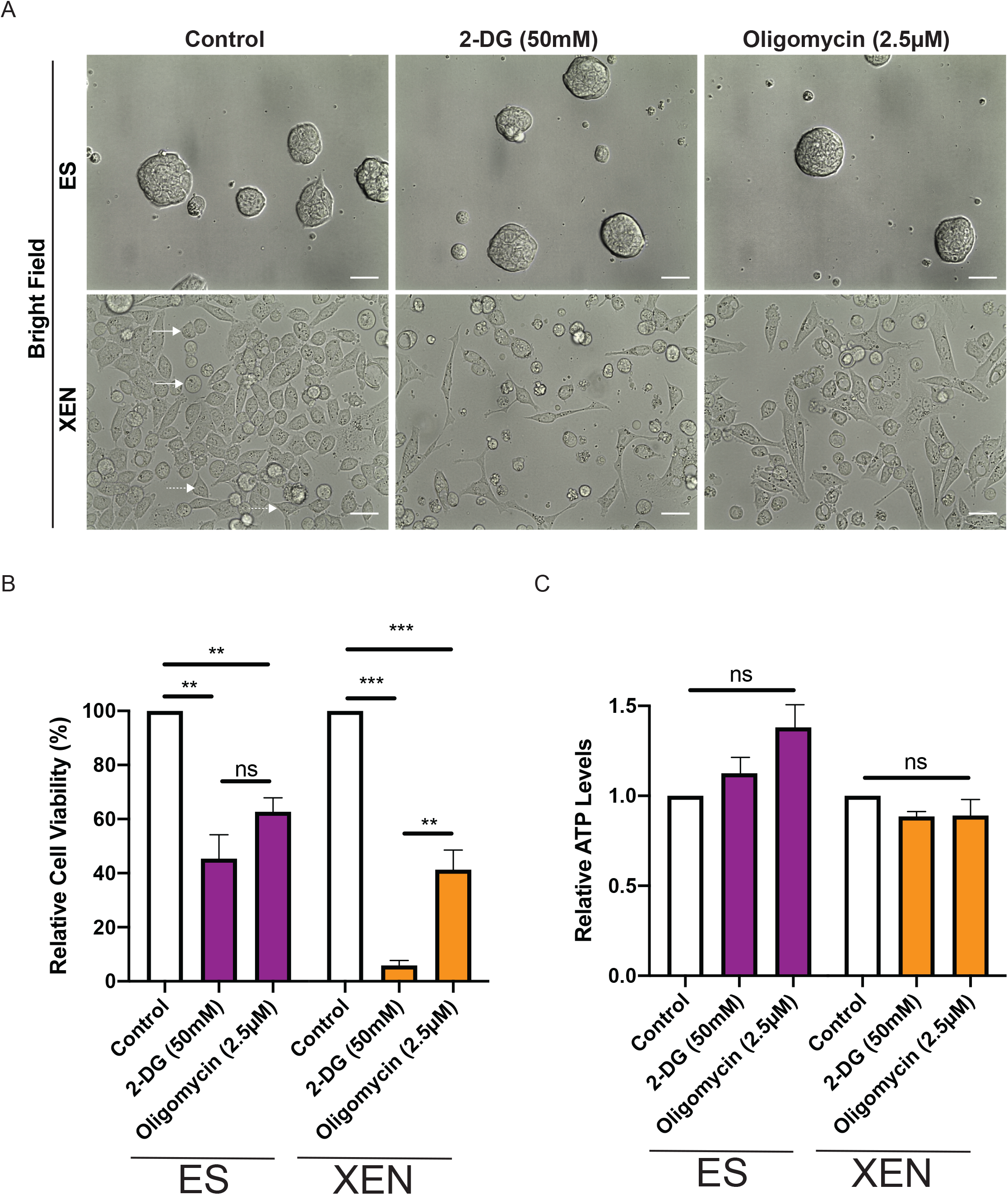
XEN cells are more sensitive to glycolytic inhibition. (A) Representative phase contrast images of ES and XEN cells cultured under control, 50mM 2-DG, and 2.5μM oligomycin. Solid arrows are indicative of round XEN morphology. Dashed arrows are indicative of stellate XEN morphology. Scale bars = 100μm. (B) Relative cell viability of ES and XEN cells under control, 50mM 2-DG and 2.5μM oligomycin conditions. (C) Total ATP levels in ES and XEN cells under control, 50mM 2-DG and 2.5μM oligomycin conditions. (n = three biological replicates, ns = not significant, \*\**P* < 0.01, \*\*\**P* <0.001).

ES colony shape appeared to be insensitive to 2-DG and oligomycin treatment; however, XEN cells lost their distinct morphologies, particularly in the 2-DG treatment and adopted a spindle-like morphology, indicative of cell stress (Figure 1A). In contrast to a previous report (Choi et al., 2020), XEN cells were more sensitive to glycolytic inhibition (5.87% ± 1.82) than OXPHOS inhibition with oligomycin (41.3% ± 7.30; *P* < 0.01; Figure 1B). Since ES cells are metabolically bivalent (Zhou et al., 2012), the significant reduction in cell viability under both treatments was expected (*P* < 0.01) and not significant between the two treatments (2-DG: 45.3% ± 8.84 *versus* oligomycin: 62.7% ± 5.24; Figure 1B). Despite this reduction in cell viability, no change in relative total ATP levels was observed between the two populations and treatments (Figure 1C). Collectively, our findings support the notion that feeder-free XEN cells are more sensitive to glycolytic inhibition than feeder-free ES cells.

### Intracellular metabolomic profiling of ICM cells

Since feeder-free ES and XEN cells responded differently to metabolic inhibitors, we exploited the feeder-free system to identify changes in intracellular metabolites between the two populations. For metabolite identification, untargeted metabolomics was used to measure the relative levels of intracellular metabolites (Figure 2). We identified 142 intracellular metabolites of which 18 were significantly enriched in ES cells and 68 in XEN cells (Table 1; Figure 2A). Next, targeted metabolomics was performed to quantify the levels of amino acids (Figure 2B) and metabolites involved in glycolysis and the tricarboxylic acid (TCA) cycle (Figure 2C). Several amino acids were significantly enriched in XEN cells, including proline, serine and threonine, which are reported to play a role in pluripotency and differentiation (Baksh et al., 2020; Comes et al., 2013; Ryu and Han, 2011; Wang et al., 2009). While most metabolites involved in glycolysis and TCA cycle were not statistically significant, intracellular lactate and 2-hydroxyglutarate were significantly higher in XEN cells when compared with ES cells (*P* < 0.05; Figure 2C). Together, our data indicate that XEN cells exhibit a unique metabolite profile that is distinct from the ES population.

**Figure 2.**
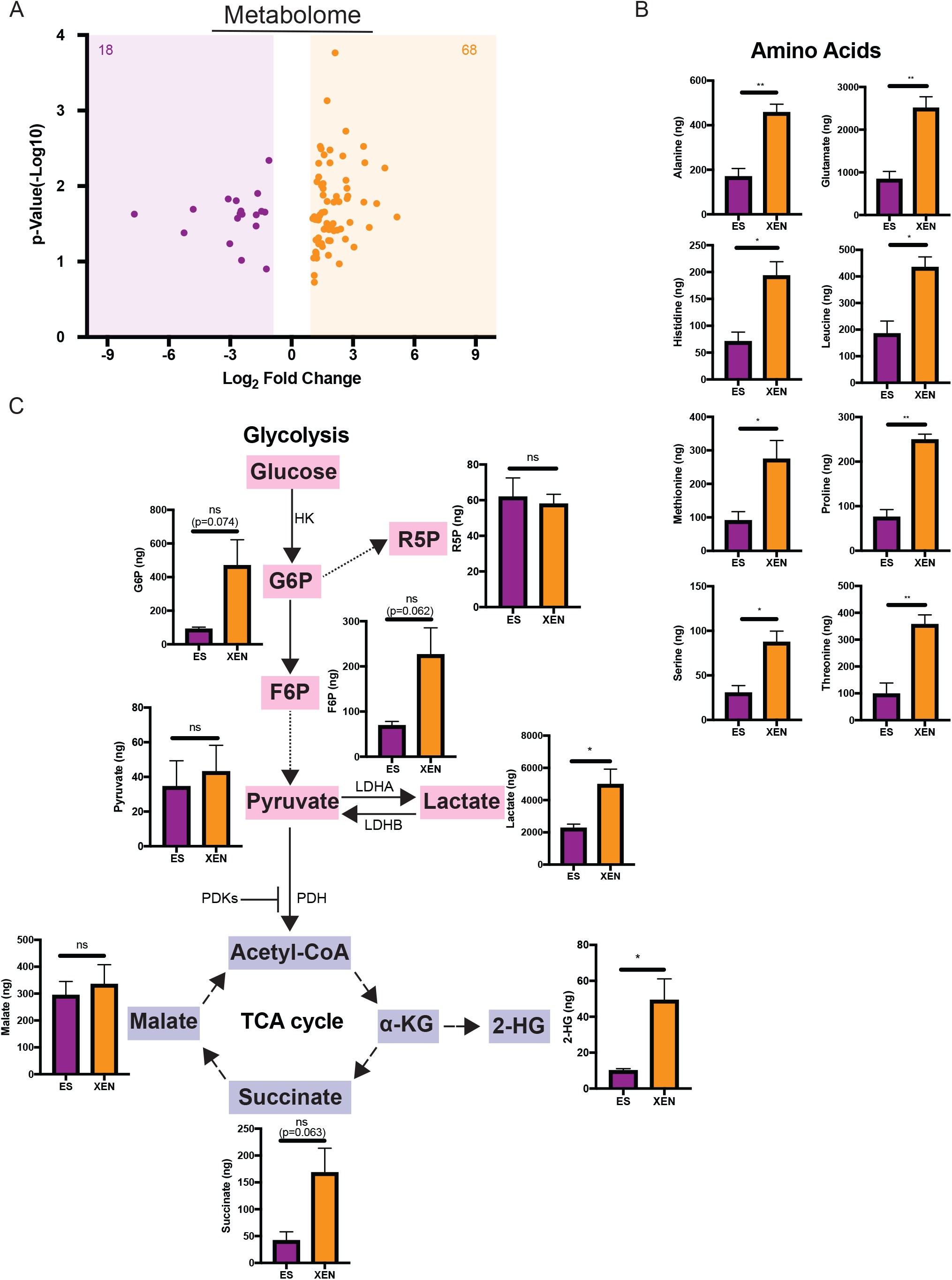
Intracellular metabolomic analysis of ES and XEN cells. (A) Volcano plots of differentially expressed metabolites from untargeted LC/GC-MS/MS metabolomics of ES and XEN cells cultured under normal conditions. (fold change > 1 and *P* < 0.05). Metabolites enriched in ES cells are highlighted in purple (38 metabolites) and metabolites enriched in XEN cells are highlighted in orange (105 metabolites). (B, C) Levels of key intracellular metabolites generated by glycolysis, TCA cycle, and amino acid detected by targeted LC/GC-MS/MS analysis of ES and XEN cells cultured under normal conditions. Metabolites are expressed in ng and were normalized to cell number. (n = three biological replicates, ns = not significant, \**P* < 0.05, \*\**P* <0.01).

**Table 1.**
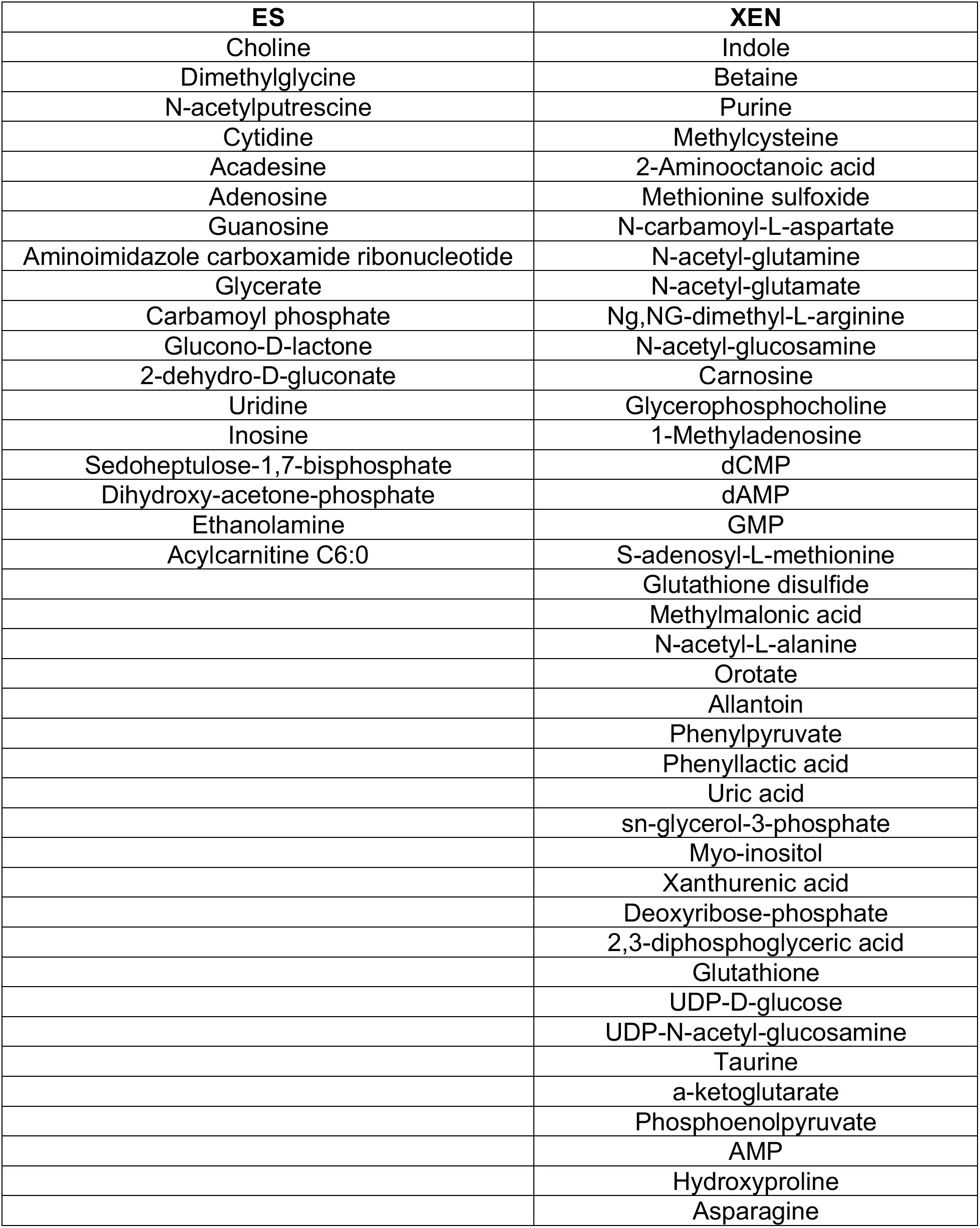
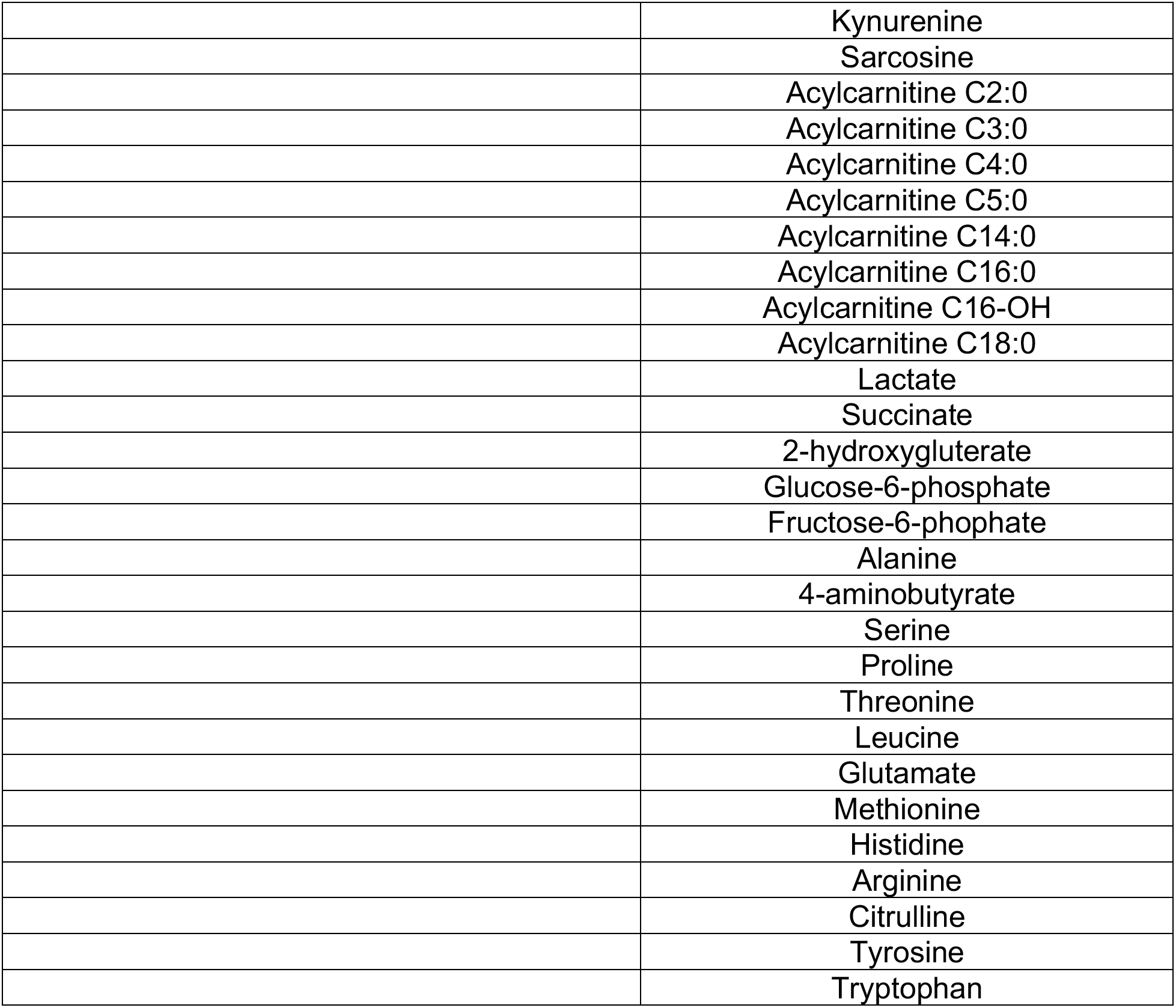
Metabolites enriched in either ES or XEN cells based on untargeted/targeted metabolomics

| ES | XEN |
| --- | --- |
| Choline | Indole |
| Dimethylglycine | Betaine |
| N-acetylputrescine | Purine |
| Cytidine | Methylcysteine |
| Acadesine | 2-Aminooctanoic acid |
| Adenosine | Methionine sulfoxide |
| Guanosine | N-carbamoyl-L-aspartate |
| Aminoimidazole carboxamide ribonucleotide | N-acetyl-glutamine |
| Glycerate | N-acetyl-glutamate |
| Carbamoyl phosphate | Ng,NG-dimethyl-L-arginine |
| Glucono-D-lactone | N-acetyl-glucosamine |
| 2-dehydro-D-gluconate | Carnosine |
| Uridine | Glycerophosphocholine |
| Inosine | 1-Methyladenosine |
| Sedoheptulose-1,7-bisphosphate | dCMP |
| Dihydroxy-acetone-phosphate | dAMP |
| Ethanolamine | GMP |
| Acylcarnitine C6:0 | S-adenosyl-L-methionine |
|  | Glutathione disulfide |
|  | Methylmalonic acid |
|  | N-acetyl-L-alanine |
|  | Orotate |
|  | Allantoin |
|  | Phenylpyruvate |
|  | Phenyllactic acid |
|  | Uric acid |
|  | sn-glycerol-3-phosphate |
|  | Myo-inositol |
|  | Xanthurenic acid |
|  | Deoxyribose-phosphate |
|  | 2,3-diphosphoglyceric acid |
|  | Glutathione |
|  | UDP-D-glucose |
|  | UDP-N-acetyl-glucosamine |
|  | Taurine |
|  | a-ketoglutarate |
|  | Phosphoenolpyruvate |
|  | AMP |
|  | Hydroxyproline |
|  | Asparagine |
|  | Kynurenine |
|  | Sarcosine |
|  | Acylcarnitine C2:0 |
|  | Acylcarnitine C3:0 |
|  | Acylcarnitine C4:0 |
|  | Acylcarnitine C5:0 |
|  | Acylcarnitine C14:0 |
|  | Acylcarnitine C16:0 |
|  | Acylcarnitine C16-OH |
|  | Acylcarnitine C18:0 |
|  | Lactate |
|  | Succinate |
|  | 2-hydroxygluterate |
|  | Glucose-6-phosphate |
|  | Fructose-6-phosphate |
|  | Alanine |
|  | 4-aminobutyrate |
|  | Serine |
|  | Proline |
|  | Threonine |
|  | Leucine |
|  | Glutamate |
|  | Methionine |
|  | Histidine |
|  | Arginine |
|  | Citrulline |
|  | Tyrosine |
|  | Tryptophan |

**Table 2.**
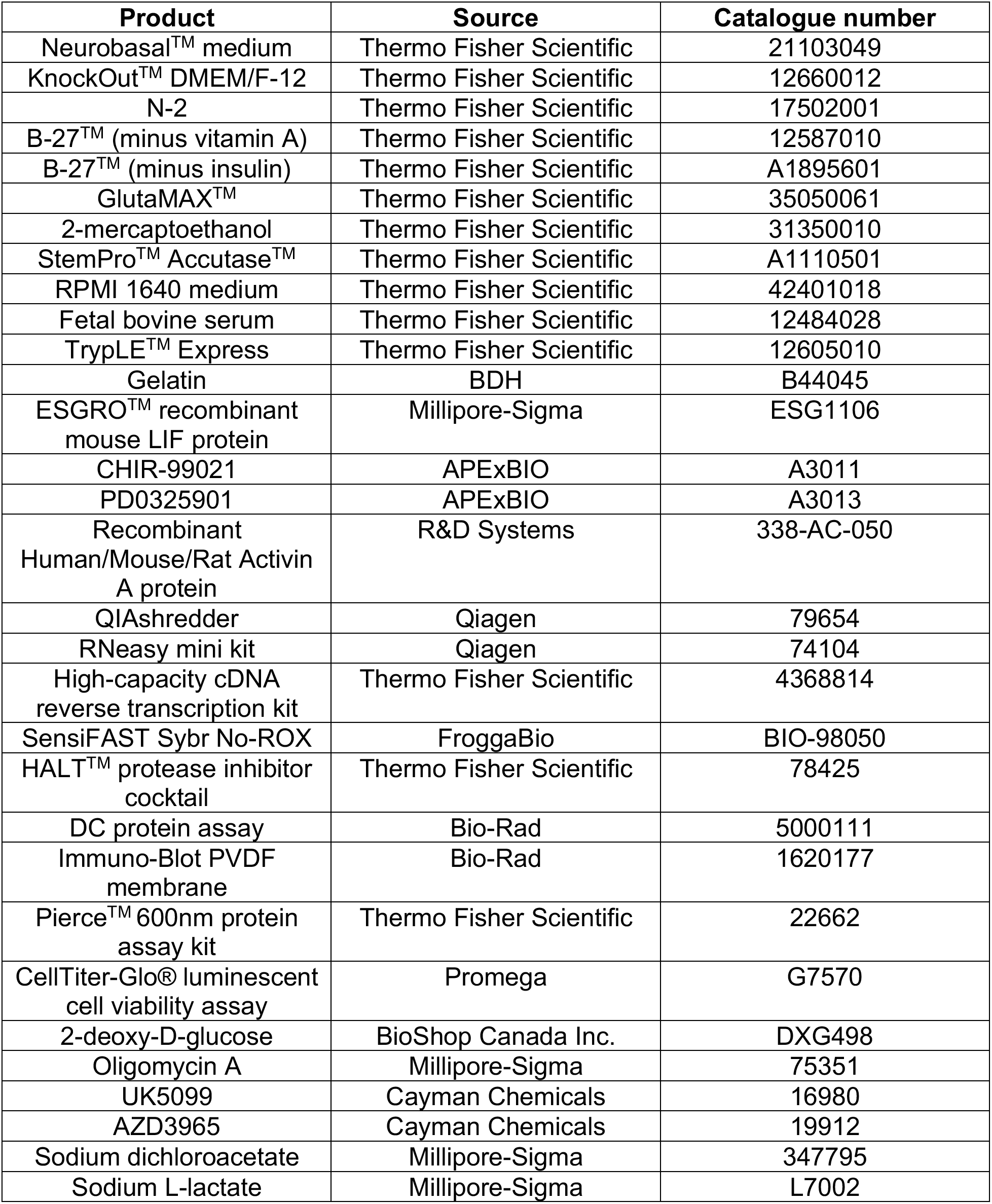
Products, manufacturer and catalogue number of items used in this report

| **Product** | **Source** | **Catalogue number** |
| --- | --- | --- |
| Neurobasal™ medium | Thermo Fisher Scientific | 21103049 |
| KnockOut™ DMEM/F-12 | Thermo Fisher Scientific | 12660012 |
| N-2 | Thermo Fisher Scientific | 17502001 |
| B-27™ (minus vitamin A) | Thermo Fisher Scientific | 12587010 |
| B-27™ (minus insulin) | Thermo Fisher Scientific | A1895601 |
| GlutaMAX™ | Thermo Fisher Scientific | 35050061 |
| 2-mercaptoethanol | Thermo Fisher Scientific | 31350010 |
| StemPro™ Accutase™ | Thermo Fisher Scientific | A1110501 |
| RPMI 1640 medium | Thermo Fisher Scientific | 42401018 |
| Fetal bovine serum | Thermo Fisher Scientific | 12484028 |
| TrypLE™ Express | Thermo Fisher Scientific | 12605010 |
| Gelatin | BDH | B44045 |
| ESGRO™ recombinant mouse LIF protein | Millipore-Sigma | ESG1106 |
| CHIR-99021 | APExBIO | A3011 |
| PD0325901 | APExBIO | A3013 |
| Recombinant Human/Mouse/Rat Activin A protein | R&D Systems | 338-AC-050 |
| QIAshredder | Qiagen | 79654 |
| RNeasy mini kit | Qiagen | 74104 |
| High-capacity cDNA reverse transcription kit | Thermo Fisher Scientific | 4368814 |
| SensiFAST Sybr No-ROX | FroggaBio | BIO-98050 |
| HALT™ protease inhibitor cocktail | Thermo Fisher Scientific | 78425 |
| DC protein assay | Bio-Rad | 5000111 |
| Immuno-Blot PVDF membrane | Bio-Rad | 1620177 |
| Pierce™ 600nm protein assay kit | Thermo Fisher Scientific | 22662 |
| CellTiter-Glo® luminescent cell viability assay | Promega | G7570 |
| 2-deoxy-D-glucose | BioShop Canada Inc. | DXG498 |
| Oligomycin A | Millipore-Sigma | 75351 |
| UK5099 | Cayman Chemicals | 16980 |
| AZD3965 | Cayman Chemicals | 19912 |
| Sodium dichloroacetate | Millipore-Sigma | 347795 |
| Sodium L-lactate | Millipore-Sigma | L7002 |

**Table 3.** Primary and secondary antibodies used in this report

| **Antibody** | **Source** | **Catalogue number** |
| --- | --- | --- |
| Mouse monoclonal to PDK1 | Santa Cruz | sc-293160 |
| Mouse monoclonal to PDK4 | Santa Cruz | sc-28783 |
| Rabbit polyclonal to PDC-E1α<sup>pSer232</sup> | Millipore-Sigma | AP1063 |
| Rabbit polyclonal to PDC-E1α<sup>pSer293</sup> | Novus Biologicals | NB110-93479 |
| Mouse monoclonal to PDC-E1α | Abcam | Ab110334 |
| Rabbit monoclonal to LDHA | Cell Signalling Technology | 3582 |
| Mouse monoclonal to LDHB | Abcam | Ab85319 |
| Mouse monoclonal to β-ACTIN | Santa Cruz | sc-69879 |
| Rabbit polyclonal to DAB2 | Cell Signalling Technology | 12906S |
| Goat polyclonal to GATA6 | Novus Biologicals | AF1700 |
| Rat monoclonal to KERATIN-8 | DSHB | TROMA-I-f |
| Rabbit polyclonal to OCT4 | Cell Signalling Technology | 2750 |
| Goat anti-rabbit IgG-HRP | Millipore-Sigma | AQ132P |
| Goat anti-mouse IgG-HRP | Millipore-Sigma | A5278 |
| Goat anti-rat IgG-HRP | Abcam | Ab97057 |
| Rabbit anti-goat IgG-GRP | Millipore-Sigma | A8919 |

**Table 4.**
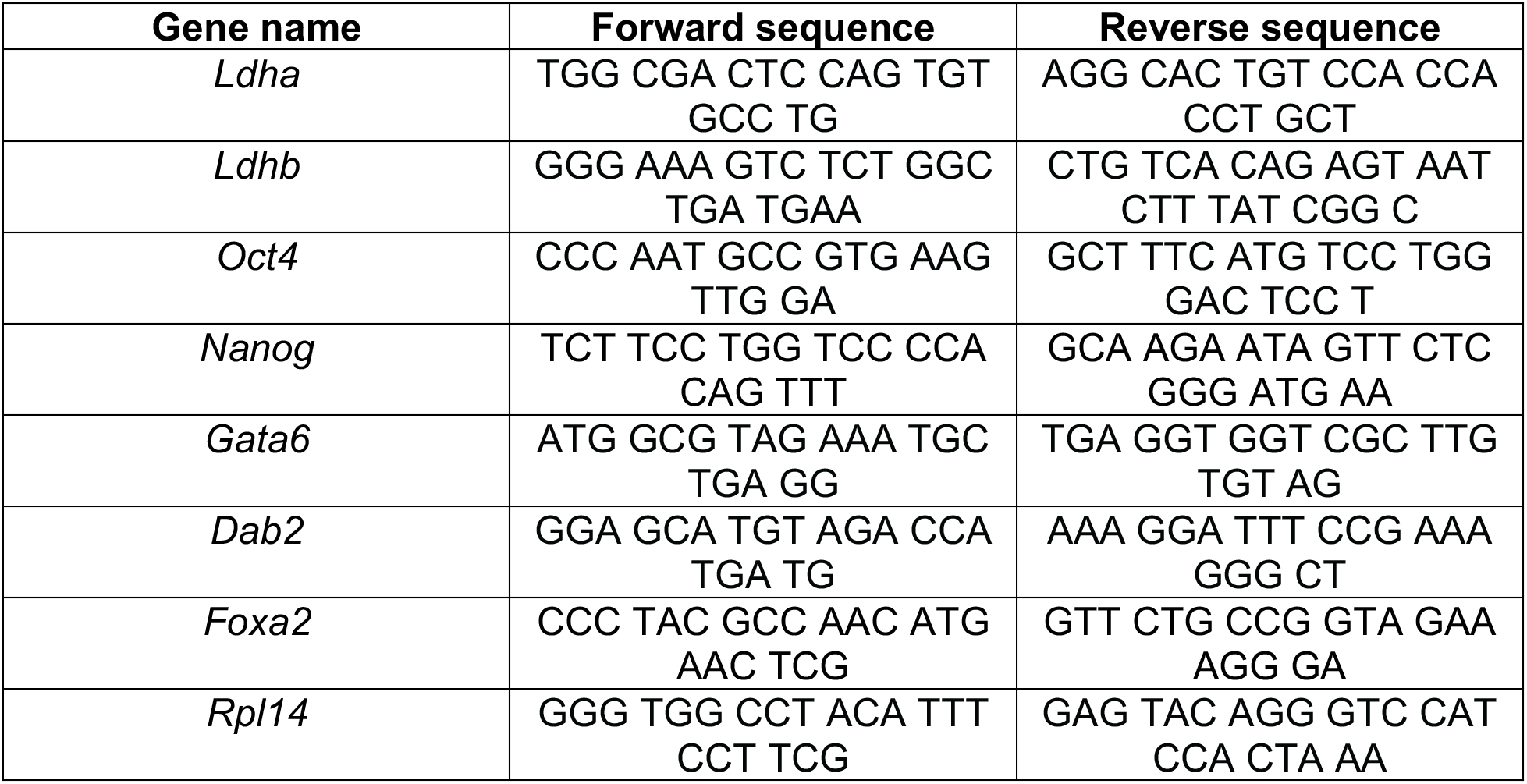
Primer sequences used in this report

### Transcriptomic and proteomic analysis showcases the importance of metabolic pathways during embryonic development

Bulk transcriptomic and proteomic analyses were performed on ES and XEN cells in order to compare global gene expression and protein abundance patterns and assign molecular pathway(s) to the metabolite profiles. Results revealed 2742 genes (Figure 3A) and 165 proteins (Figure 3B) significantly upregulated in the ES population. In contrast, 1716 genes (Figure 3A) and 298 proteins (Figure 3B) were significantly upregulated in XEN cells. As expected, markers of pluripotency including *Nanog*, *Sox2*, OCT4 and ESRRB were enriched in the ES population, while XEN markers such as *Gata4/6*, *Sox7/17*, *Dab2*, and PDGFRA were significantly enriched in XEN cells (Figure 3A, B). Differentially expressed genes and proteins enriched in either ES or XEN cells were selected and analyzed by gene ontology to classify targets based on their biological process (Figure S2A-D), cellular compartment (Figure S2E-H) and molecular process (Figure S2I-L). As expected, targets from ES cells were enriched for biological processes associated with multicellular organism development and regulation of transcription (Figure S3A, B). Conversely, XEN cells terms were associated with various signaling pathways, endodermal development and extracellular matrix organization (Figure S3C, D).

**Figure 3.**
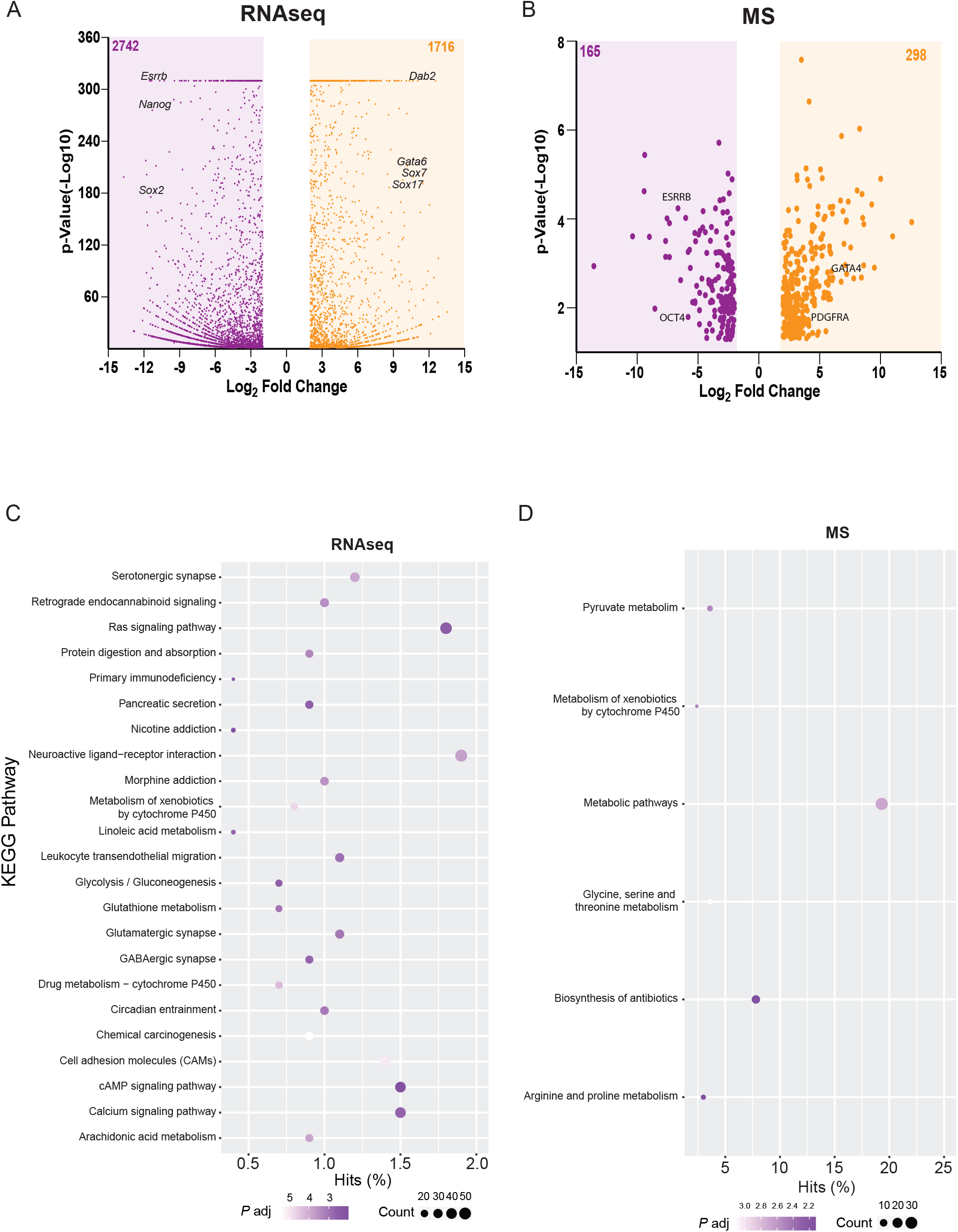
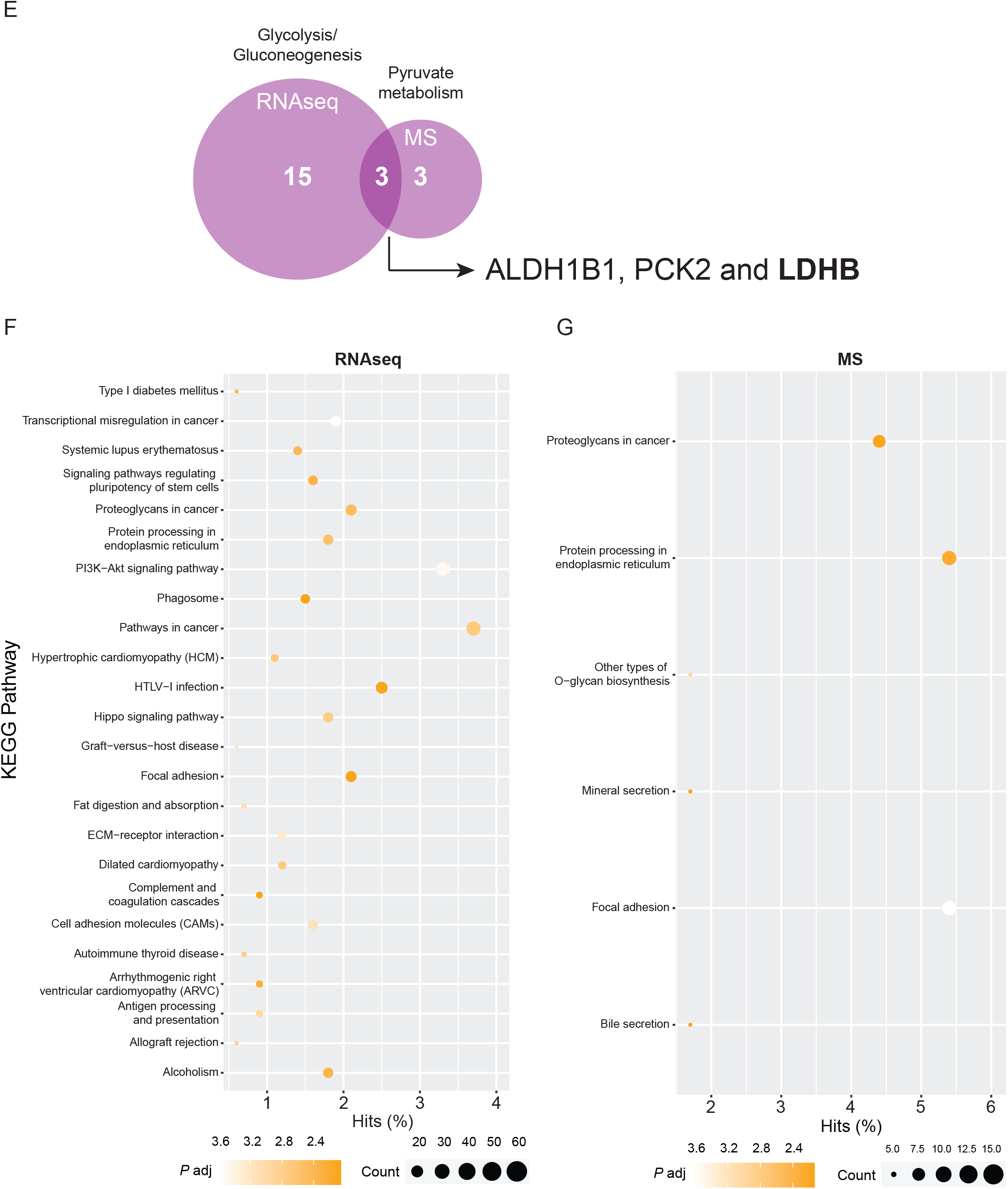
Bulk transcriptomic and proteomic analysis of ES and XEN cells. (A) Volcano plots of differentially expressed genes from bulk RNA sequencing of ES and XEN cells cultured under normal conditions. (fold change > 2 and adjusted *P* < 0.01). Genes enriched in ES cells are highlighted in purple (2742 genes) and genes enriched in XEN cells are highlighted in orange (1716 genes). (B) Volcano plots of differentially expressed proteins from bulk proteomics of ES and XEN cells cultured under normal conditions. (fold change > 2 and adjusted *P* < 0.01). Proteins abundant in ES cells are highlighted in purple (165 proteins) and proteins abundant in XEN cells are highlighted in orange (298 proteins). (C) Dot plot representing enriched KEGG pathways in ES cells from transcriptomic dataset. (D) Dot plot representing enriched KEGG pathways in ES cells from proteomic dataset. (E) Venn diagram of common targets from both transcriptomic and proteomic datasets for specific KEGG pathways. (F) Dot plot representing enriched KEGG pathways in XEN cells from transcriptomic dataset. (G) Dot plot representing enriched KEGG pathways in XEN cells from proteomic dataset.

Kyoto Encyclopedia of Genes and Genomes (KEGG) enrichment analysis was also performed on genes and proteins to identify biological pathways specific to each population (Figure 3C-G). XEN cells expressed targets primarily involved in proteoglycan metabolism and endoplasmic reticulum processing (Figure 3F, G, S3G, H), as reported previously (Choi et al., 2020). However, metabolism of xenobiotics by cytochrome P450 and glucose/pyruvate metabolism were enriched in the ES population and represent two common KEGG pathways between the RNAseq and MS datasets (Figure 3C, D). Interestingly, three targets that were highly enriched in both datasets and linked to glucose metabolism were aldehyde dehydrogenase 1 family member B1 (ALDH1B1), phosphoenolpyruvate carboxykinase 2 (PCK2) and lactate dehydrogenase B (LDHB, Figure 3E). Together, this data would suggest that these targets and their metabolites play a role in either maintaining pluripotency or in the differentiation towards the XEN lineage.

### Enzymes involved in lactate homeostasis are upregulated in feeder-free XEN cells

Pyruvate is converted into lactate by the activity of LDHA and converted back by LDHB; since the latter was enriched in the ES population (Figure 3A-E), and lactate was enriched in XEN cells intracellularly (Figure 2C), we sought to further investigate the levels of enzymes involved in lactate metabolism. qRT-PCR results showed significantly higher *Ldha*/*Ldhb* expression in XEN cells compared with ES cells (*P* < 0.01, Figure 4A), and this was confirmed at the protein level using immunoblot analysis (Figure 4B). Extracellular lactate levels were also measured, and results showed significantly higher levels in XEN cells (*P* < 0.05, Figure 4C). Furthermore, the expression of the lactate transporter *Mct1* (Figure 4D), but not *Mct4* (Figure 4E), was significantly upregulated in XEN cells (*P* < 0.0001) and is a prime candidate linked to these high extracellular lactate levels. Pyruvate can also be oxidized in the mitochondria by the pyruvate dehydrogenase complex (PDC) to generate acetyl-CoA. This conversion is dependent on the activity of the PDC, which is negatively regulated by phosphorylation of the E1α subunit by members of the pyruvate dehydrogenase kinase (PDK) family. Expression analysis shows that *Pdk1, 2* and *4,* but not *Pdk3,* were significantly downregulated in XEN cells (Figure 4F-I). Similarly, immunoblot analysis revealed low levels of PDC-E1α^Ser232/Ser293^ in XEN cells (Figure 4J), indicating that the PDC is active and would suggest higher OXPHOS activity. However, expression analysis of the mitochondrial pyruvate carriers *Mpc1* (Figure 4K), but not *Mpc2* (Figure 4L), showed significantly reduced expression in XEN cells (*P* < 0.0001). Thus, despite having an active PDC, pyruvate in XEN cells fails to enter the mitochondria, and instead is preferentially converted to lactate by the enhanced LDHA activity (Figure 4A, B). These results would suggest that lactate participates in specifying ES cells towards a XEN lineage.

**Figure 4.**
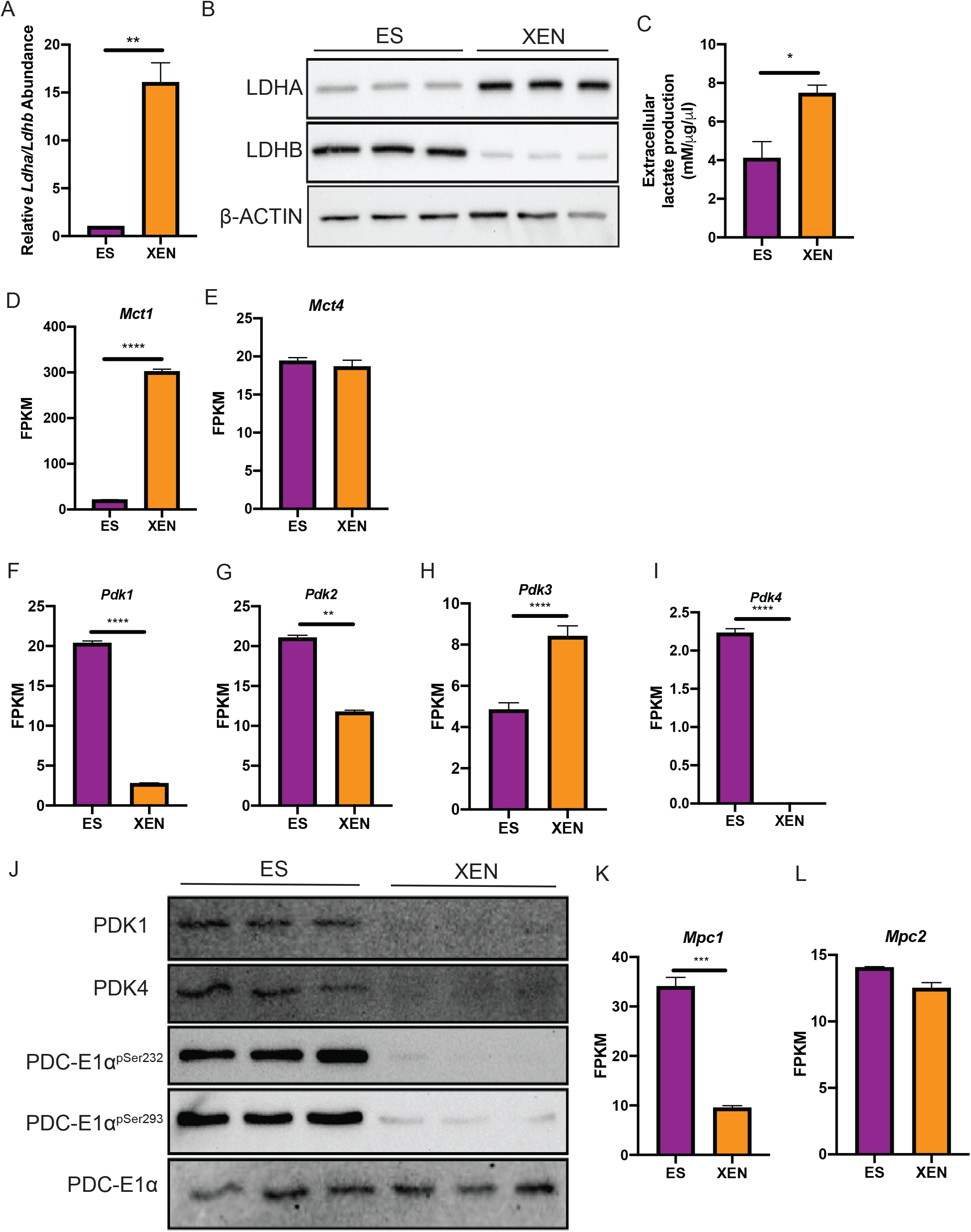
Levels of enzymes involved in pyruvate metabolism in ES and XEN cells. (A) Bar graph of qRT-PCR analysis of *Ldha/Ldhb* ratio in ES and XEN cells. *Rpl14* was used a constitutive gene for qRT-PCR. (n = three biological replicates, \*\**P* <0.01). (B) Representative immunoblot analysis showing levels of LDHA and LDHB in ES and XEN cells. β-ACTIN served as a loading control. (n = three biological replicates). (C) Bar graph of extracellular quantification of lactate levels in ES and XEN cells by BioProfile^®^400 Chemical Analyzer. (n = three biological replicates, \**P* < 0.05). (D-E) Bar graph of expression level of *Mct1* and *Mct4* in ES and XEN cells. FPKM was used for RNAseq data. (n = three biological replicates, \*\*\*\**P* <0.0001). (F-I) Bar graph of FPKM of *Pdk1-4* in ES and XEN cells. (n = three biological replicates, ns = not significant, \*\**P* <0.01, \*\*\*\**P* <0.0001). (J) Representative immunoblot analysis showing levels of PDK1, PDK4, PDC-E1α^pSer232^, PDC-E1α^pSer293^, and PDC-E1α in ES and XEN cells. (n = three biological replicates). (K-L) Bar graph of expression level of *Mpc1* and *Mpc2* in ES and XEN cells. FPKM was used for RNAseq data. (n = three biological replicates, \*\*\**P* <0.001).

### Promoting intracellular lactate enhances XEN differentiation

Given the previous data highlighting a potential role for lactate in XEN cells (Figure 1–4), we sought to directly assess whether modulating lactate metabolism had an effect on XEN induction *in vitro* (Figure 5). We adopted a previously defined chemical cocktail (Anderson et al., 2017) to induce naïve mouse ES cells towards the XEN lineage. Exogenous supplementation of LIF, Activin A and CHIR99021 (inhibitor of glycogen synthase kinase 3) in the absence of insulin (Figure S3A) promotes differentiation towards the XEN lineage as evident by elevated *Gata6* and *Dab2* transcript and low levels of *Nanog* and *Oct4* (Figure S3B-E). To test whether lactate metabolism plays a role in XEN differentiation, we used two approaches. In the first, three inhibitors were selected, two of which would promote intracellular lactate accumulation (UK5099 and AZD3965), while the other would promote pyruvate uptake into the mitochondria (Dichloroacetate, DCA). In the second approach, media was supplemented with exogenous L-lactate and cells were assessed for markers of pluripotency and XEN differentiation. UK5099 is a potent pan MPC inhibitor that prevents pyruvate uptake into the mitochondria (Halestrap, 1975; Vacanti et al., 2014), while AZD3965, a highly-specific MCT1 inhibitor, reduces lactate secretion in human stem cells (Gu et al., 2016) and cancer cells (Polanski et al., 2014). The latter was used since MCT1 is the predominant transporter responsible for exporting lactate extracellularly in XEN cells (Figure 4D). Generally, neither UK5099, AZD3965 or L-lactate affected *Oct* or *Nanog* expression (Figure 5C, F, G, J, K); however, *Oct4* expression was slightly but significantly downregulated (*P* < 0.05) in ES cells cultured under 2i-LIF and UK5099 conditions (Figure 5B). In contrast, DCA-treated ES cells cultured with 2i-LIF had slightly elevated *Oct4* and *Nanog* expression when compared to controls (Figure 5N, O).

**Figure 5.**
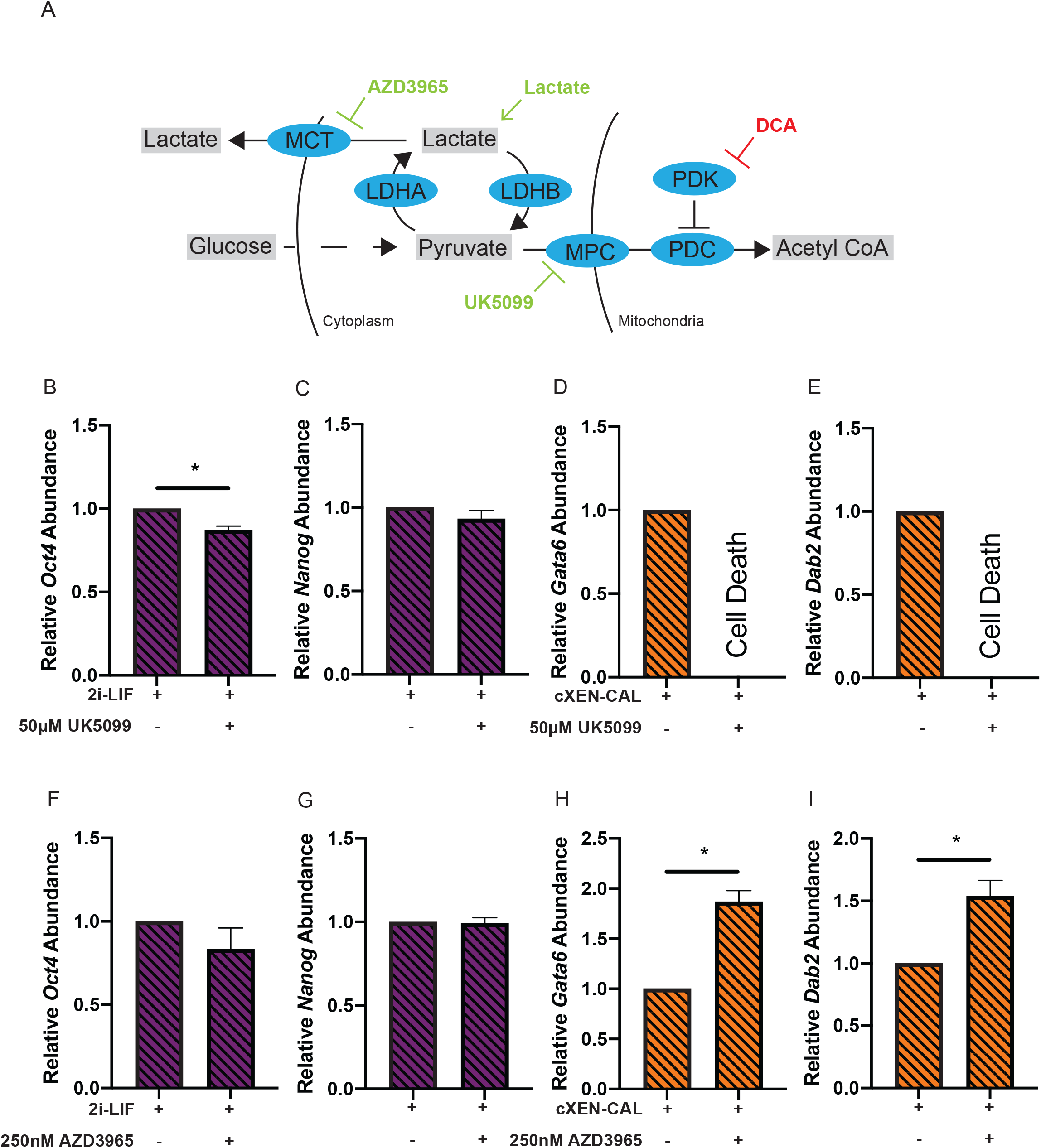
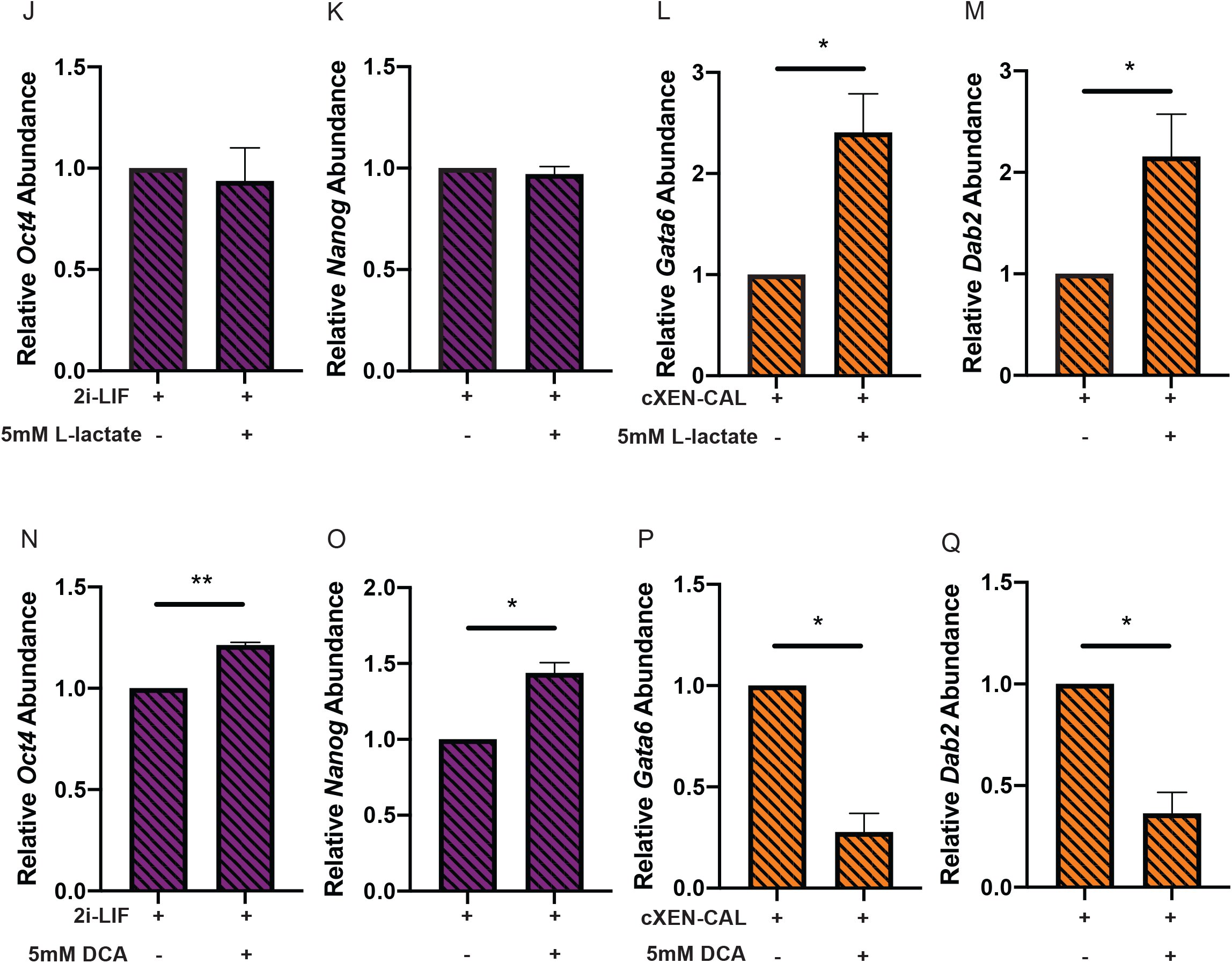
Intracellular lactate enhances XEN differentiation in vitro. (A) Schematic diagram of glucose metabolism and the metabolic enzymes involved. Green is indicative of treatments that increase intracellular lactate and red is indicative of treatment that decrease intracellular lactate. (B-C) Bar graph of expression level of *Oct4* and *Nanog* in ES cells in 2i-LIF with or without 50μM UK5099 for 48h. *Rpl14* was used a constitutive gene for qRT-PCR. (n = three biological replicates, \**P* <0.05). (D-E) Bar graph of expression level of *Gata6* and *Dab2* in ES cells induced towards a XEN lineage with or without 50μM UK5099 for 96h. *Rpl14* was used a constitutive gene for qRT-PCR. (n = three biological replicates, \**P* <0.05). (F-G) Bar graph of expression level of *Oct4* and *Nanog* in ES cells in 2i-LIF with or without 250nM AZD3965 for 48h. *Rpl14* was used a constitutive gene for qRT-PCR. (n = three biological replicates). (H-I) Bar graph of expression level of *Gata6* and *Dab2* in ES cells induced towards a XEN lineage with or without 250nM AZD3965 for 96h. *Rpl14* was used a constitutive gene for qRT-PCR. (n = three biological replicates, \**P* <0.05). (J-K) Bar graph of expression level of *Oct4* and *Nanog* in ES cells in 2i-LIF with or without 5mM L-lactate for 48h. *Rpl14* was used a constitutive gene for qRT-PCR. (n = three biological replicates). (L-M) Bar graph of expression level of *Gata6* and *Dab2* in ES cells induced towards a XEN lineage with or without 5mM L-lactate for 96h. *Rpl14* was used a constitutive gene for qRT-PCR. (n = three biological replicates, \**P* <0.05). (N-O) Bar graph of expression level of *Oct4* and *Nanog* in ES cells in 2i-LIF with or without 5mM DCA for 48h. *Rpl14* was used a constitutive gene for qRT-PCR. (n = three biological replicates, \**P* <0.05, \*\**P* <0.01). (P-Q) Bar graph of expression level of *Gata6* and *Dab2* in ES cells induced towards a XEN lineage with or without 5mM DCA for 96h. *Rpl14* was used a constitutive gene for qRT-PCR. (n = three biological replicates, \**P* <0.05).

Next, ES cells were induced towards the XEN lineage under control conditions or treated with inhibitors or supplemented with L-lactate. ES cells treated with UK5099 in XEN induction media were not viable after 48h post treatment (Figure 5D, E), while surprisingly those cultured with either AZD3965 (Figure 5H, I) or supplemented with L-lactate (Figure 5L, M) for 96h showed enhanced *Gata6* and *Dab2* expression when compared to controls. Prolonged L-lactate supplementation induced cell death at 10 days post treatment in cells differentiated towards the XEN lineage (data not shown). Conversely, when ES cells were treated with DCA in XEN induction media, thereby promoting pyruvate uptake into the mitochondria and reducing intracellular lactate, *Gata6* and *Dab2* expression were significantly downregulated when compared to controls (*P* < 0.05, Figure 5P, Q). Therefore, promoting intracellular lactate accumulation enhanced XEN induction *in vitro*, highlighting a novel role for lactate as a signaling molecule involved in the differentiation of XEN cells.

## Discussion

We have shown previously that F9 embryonal carcinoma cells mimic the transition to a XEN-like lineage when co-treated with retinoic acid and a cAMP analog, and more importantly, these cells exhibited a metabolic phenotype reliant exclusively on glycolysis (Gatie and Kelly, 2018). While recent reports have expanded on the metabolic profiles observed in the early embryonic lineages (Choi et al., 2020; Zhou et al., 2012), a comprehensive analysis of the metabolites and their roles during development remains lacking. Our data indicates that feeder-free XEN cells are sensitive to glycolytic inhibition when compared with OXPHOS inhibition (Figure 1A, B). This phenomenon is also observed in mouse primed stem cells, human stem cells (Zhou et al., 2012) and canine epiblast stem cells (Tobias et al., 2018), suggesting that these characteristics are adaptive mechanisms linked to a developmental stage. Furthermore, XEN cells possess altered glycolytic enzyme levels involved in pyruvate and lactate homeostasis (Figure 4). Recent reports suggested that XEN cells rely on OXPHOS metabolism for energetic needs (Choi et al., 2020); however, the developmental stage and culture condition of XEN cells may influence their metabolic profile. For instance, primitive endoderm cells are present within the ICM at 3.5 days post fertilization and they further commit into parietal and visceral endoderm cells shortly after implantation. A similar phenomenon is observed between preimplantation naïve *versus* post-implantation primed ES cells, which display distinct metabolic profiles (Zhou et al., 2012), suggesting that XEN cells may exhibit a unique metabolic signature depending on their developmental stage. In addition, the localization of XEN cells varies depending on the stage of development, where primitive endoderm cells reside within the ICM, visceral endoderm cells surround the epiblast while parietal endoderm cells interact with trophoblast stem cells and undergo an epithelial-to-mesenchymal transition and migrate along the inner surface of the trophectoderm (Hogan et al., 1980). Therefore, the precise identity of these XEN cells and their interaction with the microenvironment may influence their metabolic needs. Lastly, culture conditions vary between trophoblast, ES and XEN cells, and its known that the latter can be derived using various culture conditions and supplements (Anderson et al., 2017; Cho et al., 2012; Fujikura et al., 2002; He et al., 2020; Kunath et al., 2005; McDonald et al., 2014; Niakan et al., 2013; Paca et al., 2012; Shimosato et al., 2007; Soprano et al., 2007; Wamaitha et al., 2015). Together, the ability to derive XEN cells by various means would contribute to their heterogeneity (Kunath et al., 2005; Paca et al., 2012) and potentially affect the metabolic profile of these cells. For example, XEN cells can be generated by the overexpression of *Gata4* in ES cells (Fujikura et al., 2002; Wamaitha et al., 2015) or reprogramming fibroblasts using a chemical cocktail (He et al., 2020). While both methods result in the establishment of XEN cells, *Gata4*-induced ES cells display gradual reduction in hexokinase 2 (HK2) and glucose transporter 1 (GLUT1) levels (Mulvey et al., 2015), while XEN cells reprogrammed from fibroblasts display elevated *Hk2* and *Glut1* levels (He et al., 2020). Together, these differences highlight the metabolic complexity and plasticity of all cells in the developing embryo, and in XEN cells specifically.

As the embryo develops its metabolic needs shift to meet energetic demands for growth and development and this metabolic rewiring is dependent on extrinsic and intrinsic cues. The blastocyst resides in a hypoxic environment, with oxygen tension dropping significantly during implantation (Fischer and Bavister, 1993). This hypoxic environment results in the stabilization of hypoxia-inducible factor one alpha (HIF1α), resulting in the increase in enzymes shifting the metabolic profile towards glycolysis. During implantation, however, angiogenesis promotes reoxygenation of the embryo triggering a switch to OXPHOS. Despite residing in a similar environment, trophoblast stem cells rely on OXPHOS metabolism, largely to fuel the sodium/potassium ion pumps required for blastocoel expansion (Houghton et al., 2003). While extrinsic factors such as the microenvironment are important, intrinsic factors play a key role in remodeling metabolic needs to meet developmental programs. For example, naïve ES cells are metabolically flexible and utilize both glycolysis and OXPHOS metabolism. However, the transition to a primed state is marked by a switch to glycolytic metabolism, which is largely due to HIF1α stabilization within the cell (Zhou et al., 2012). While changes in the levels of metabolic enzymes play a role in remodelling cellular metabolism, the metabolites resulting from the activity of these enzymes are instrumental in regulating pluripotency and/or differentiation. Our study is the first to provide a comprehensive snapshot of the intracellular metabolite profile of feeder-free XEN cells and reveals how elevated levels of key metabolites, mainly lactate, influence differentiation.

Lactate, the primary source of carbon for the TCA cycle in both normal and cancer tissues (Hui et al., 2017), is generated by the activity of LDHA, reportedly enriched in XEN cells (Figure 4A, B). Since lactate can be converted to pyruvate by LDHB, the stoichiometric ratio between the two isoforms dictates the fate of pyruvate and lactate. In addition, the oxidation of pyruvate by the PDC is required for its conversion to acetyl CoA, and the activity of the complex is regulated by the PDK family. Despite detecting elevated intracellular (Figure 2C) and extracellular (Figure 4C) lactate levels in XEN cells, the PDC was not phosphorylated and is therefore active in XEN cells (Figure 4J). While reduced expression of *Mpc1* was found in XEN cells (Figure 4K), which may explain the rerouting of pyruvate to lactate, PDC is known to have moonlighting abilities and can translocate to the nucleus where it generates acetyl CoA for acetylation of H3K9 and H3K18 (Sutendra et al., 2014). Thus, since total lysates were used to analyze the phosphorylation status of the PDC in ES and XEN cells, it is possible that a higher proportion of the active complex is nuclear and not mitochondrial. Nevertheless, since elevated intracellular lactate levels were found in XEN cells (Figure 2C), it was hypothesized that lactate plays a crucial role during XEN differentiation. Chemical inhibition to promote intracellular lactate accumulation (Figure 5H, I) or supplementation of XEN induction media with L-lactate (Figure 5L, M) showed enhanced XEN differentiation. Surprisingly, blocking pyruvate uptake into the mitochondria using UK5099 led to cell mortality (Figure 5D, E). Alternatively, promoting pyruvate uptake into the mitochondria by inhibiting PDKs resulted in significant reduction in XEN differentiation (Figure 5P, Q). Therefore, our results support the notion that maintaining intracellular lactate levels potentiates ES cells towards a XEN lineage. Since lactate can stabilize HIF1α and promote a glycolytic phenotype (De Saedeleer et al., 2012), the lactate-HIF1α axis may provide a mechanism for XEN differentiation. In addition, lactate can also inhibit histone deacetylases (Latham et al., 2012), and in conjunction with active PDC (Figure 4J), promote hyperacetylation, which we observed in XEN cells (data not shown). Thus, it appears lactate is a key player linking metabolism with epigenetic factors and transcription. These links provide insight into the role lactate plays in directing the differentiation of XEN cells, and together they underscore the importance of how the metabolic profile directs lineage commitment within the mouse early embryo.

## Experimental Procedures

### ES, XEN and cXEN Cell Culture

The ES-E14TG2a embryonic stem (ES) cells were obtained from the University of California, Davis. ES cells were cultured feeder-free on 0.1% gelatin-coated tissue culture plates in ES media: 50/50 neurobasal/DMEM-F12 media (Thermo Fisher), 1X N2 supplement (Thermo Fisher), 0.5X B27 (without retinoic acid; Thermo Fisher), 2mM GlutaMAX™ (Thermo Fisher), 0.1mM β-mercaptoethanol (Thermo Fisher), 100U/ml units of LIF (Millipore-Sigma), 3μM CHIR99021 (ApexBio) and 1μM PD 0325901 (ApexBio). ES cells were passaged every 3–4 days using Accutase (Thermo Fisher) and used to passage 30. ES media was changed daily to maintain optimal growth and reduce spontaneous differentiation. E4 extraembryonic endoderm (XEN) cells, generously donated from Dr. Janet Rossant, University of Toronto (Kunath et al., 2005), were cultured in RPMI1640 media (Thermo Fisher) supplemented with 15% fetal bovine serum (Thermo Fisher), 2mM GlutaMAX™ and 0.1mM β-mercaptoethanol. XEN cells were passaged every 2–4 days using TrypLE Express (Thermo Fisher) up to passage 20.

Chemically induced XEN (cXEN) differentiation was performed following a previously established protocol (Anderson et al., 2017). Briefly, ES cells were seeded onto 0.1% gelatin-coated tissue culture plates in base XEN media (RPMI1640 media, 0.5X B27 without insulin, 2mM GlutaMAX™ and 0.1mM β-mercaptoethanol) for 2 days. To induce XEN differentiation, base XEN media was supplemented with 100U/ml units of LIF, 3μM CHIR99021 and 20ng/ml Activin A (R&D Systems), which was changed every 2 days. All lines were grown at 37°C and 5% CO2 and checked for chromosomal abnormalities and mycoplasma (Dobrovolny and Bess, 2011) at the beginning of the project.

### RNA Isolation, cDNA synthesis and qRT-PCR analysis

RNA was extracted from cells using QIAshredder/RNAeasy Mini kit (Qiagen) and reverse transcribed into cDNA using the High-Capacity cDNA Reverse Transcription kit (Thermo Fisher). For quantitative RT-PCR, reactions contained 500nM of forward and reverse primers (Gatie and Kelly, 2018), SensiFAST SYBR Mix (FroggaBio), and cDNA. Reactions were run on a CFX Connect Real-Time PCR detection system (Bio-Rad), and results presented using the comparative cycle threshold (2^−ΔΔCt^) method with *Rpl14* serving as the internal control.

### RNA Sequencing Analysis

Total RNA was extracted as described previously and quantified using a NanoDrop 2000 spectrophotometer (Thermo Fisher) and Agilent 2100 bioanalyzer (Agilent Technologies). Library construction and sequencing were performed using the BGISEQ-500 platform (Beijing Genome Institute). Clean reads were mapped to a reference genome using Bowtie2 (Langmead and Salzberg, 2012) and gene expression levels were calculated with RSEM (Li and Dewey, 2011). Differentially expressed genes were detected with DEseq2 (Love et al., 2014) with fold change ≥ 2.0 and *P* ≤ 0.0001, and GO-term enrichment and KEGG analyses were conducted using David v6.8 (Huang da et al., 2009a, b).

### Protein extraction, quantification and immunoblot analysis

Total cell lysates were harvested using RIPA buffer supplemented with protease and phosphatase inhibitors (Thermo Fisher). Protein concentrations were quantified using a DC™ protein assay (Bio-Rad). Approximately 5–20μg of protein was separated on 5–15% polyacrylamide gels at 100V, then transferred onto PVDF membranes (Bio-Rad) overnight at 4°C at 20V. Membranes were washed in TBS-T with 0.1% Tween-20 and blocked with 5% skim milk powder for 30 minutes at room temperature with gentle shaking. Membranes were probed with primary antibodies (Gatie and Kelly, 2018) overnight at 4°C followed by washes with TBS-T and secondary antibody incubation for 2 hours at room temperature. Images were captured using a ChemiDoc™ Touch Imaging System (Bio-Rad).

### Proteomic Analysis

Sample preparation, handling and quantification were performed according to (Cooper et al., 2018). Briefly, cells were pelleted and 50μg of protein was quantified by ionic detergent compatible Pierce 660nm Protein Assay Reagent (Thermo Fisher) in 8M Urea, 50mM ammonium bicarbonate, 10mM dithiothreitol, and 2% SDS lysis buffer. Lysates were sonicated, reduced and alkylated followed by precipitation in chloroform/methanol (Wessel and Flugge, 1984), and digested samples were resuspended in 0.1% formic acid in preparation for LC-MS/MS. One microgram of tryptic peptides was injected into a Waters nano-Acquity HPLC system (Waters, Milford, MA) coupled to an ESI Orbitrap mass spectrometer (Orbitrap Elite or QExactive, ThermoFisher Scientific). MS raw files were examined using MaxQuant (1.6.5.0) and the Human Uniprot database. Bioinformatic analyses were performed in Perseus (1.5.8.5), and statistical analyses performed using multiple sample t-test with a permutation FDR set at 0.05. GO-term enrichment and KEGG analyses were conducted using David v6.8 (Huang da et al., 2009a, b).

### Detection of Total ATP levels

Total ATP levels were measured using the CellTiter-Glo^®^ Luminescent Cell Viability Assay (Promega). Cells were detached and re-suspended in media and aliquoted into 96-well plate. Cells were lysed in CellTiter-Glo^®^ reagent, incubated in the dark for 10 minutes, and luminance recorded using a Modulus™ II microplate multimode system (Promega).

### Detection of Extracellular Lactate Levels

Cells were cultured under normal conditions until reaching 70% confluency. Media from each cell population was collected, centrifuge at 14000g at 4°C for 10 minutes and analyzed using a BioProfile^®^400 Chemical Analyzer (Nova Biochemical), at the GCRC Metabolomics Core Facility, McGill University. Values were normalized to protein concentrations.

### Metabolomic Analysis

Three million cells, cultured for 4 days under normal conditions, per sample were used for extraction and analysis. Samples were analyzed for metabolites involved in glycolysis, and TCA cycle, as well as amino acids, fatty acid and lipids. Sample preparation, handling and quantification were carried out as in (Yuan et al., 2012) by The Analytical Facility for Bioactive Molecules, The Hospital for Sick Children, Toronto, Canada. Briefly, cells were harvested in 80% methanol (vol/vol) solution, centrifuged at 14,000g for 5 minutes at 4°C. Samples were aliquoted and lyophilized prior to injection. Approximately 20μl of LC/MS grade water was used to resuspend samples and 5–10μl was injected into a LC-MS/MS system which was a SCIEX 5500 QTRAP mass spectrometer (SCIEX) coupled with an Agilent 1290 HPLC stack (Agilent Technologies). MultiQuant (v3.0; SCIEX) was used for analysis,

## Data and Code Availability

Data files can be obtained from accession number GEO: GSE159855.

## Statistical Analysis

All values are presented as mean ± SEM from at least three biological experiments. In some instances (TEM experiments), three technical replicates were also included. Comparisons between two groups were performed using Student’s t-test, while comparisons between three or more groups were done using an ANOVA test followed by Tukey’s honest significant difference test. All graphs and statistics were generated using Prism (v8.4.3). *P*-values were considered significant at ^*^*P*<0.05, ^**^*P*<0.01, ^***^*P*<0.001, ^****^*P*<0.0001.

## Acknowledgments

This work was supported by discovery grant from the Natural Sciences and Engineering Research Council (NSERC R2615A02) to G.M.K. G.A.L. was supported by NSERC and Canadian Foundation for Innovation (CFI). M.I.G was supported by NSERC CGS-D scholarship and the Child Health Research Institute. The authors acknowledge the assistance from the GCRC Metabolomics core facilities, which is supported by grants from CFI, Canadian Institutes of Health Research, and Terry Fox Research Institute. The authors wish to thank Ashley St. Pierre of The Analytical Facility for Bioactive Molecules, The Hospital for Sick Children, Toronto, Canada for assistance with metabolite identification and analysis. The authors would like to thank Rached Alkallas (McGill) for assistance with RNA sequencing and data visualization.

## Author Contributions

MIG conceptualized, designed, performed the experiments and wrote the manuscript.

TTC performed and analyzed proteomics experiment.

GAL managed the proteomic component of the project.

GMK was involved with MIG and the experimental design and supervised the research.

## Declaration of Interests

The authors declare no interest.

---

**Figure S1.**
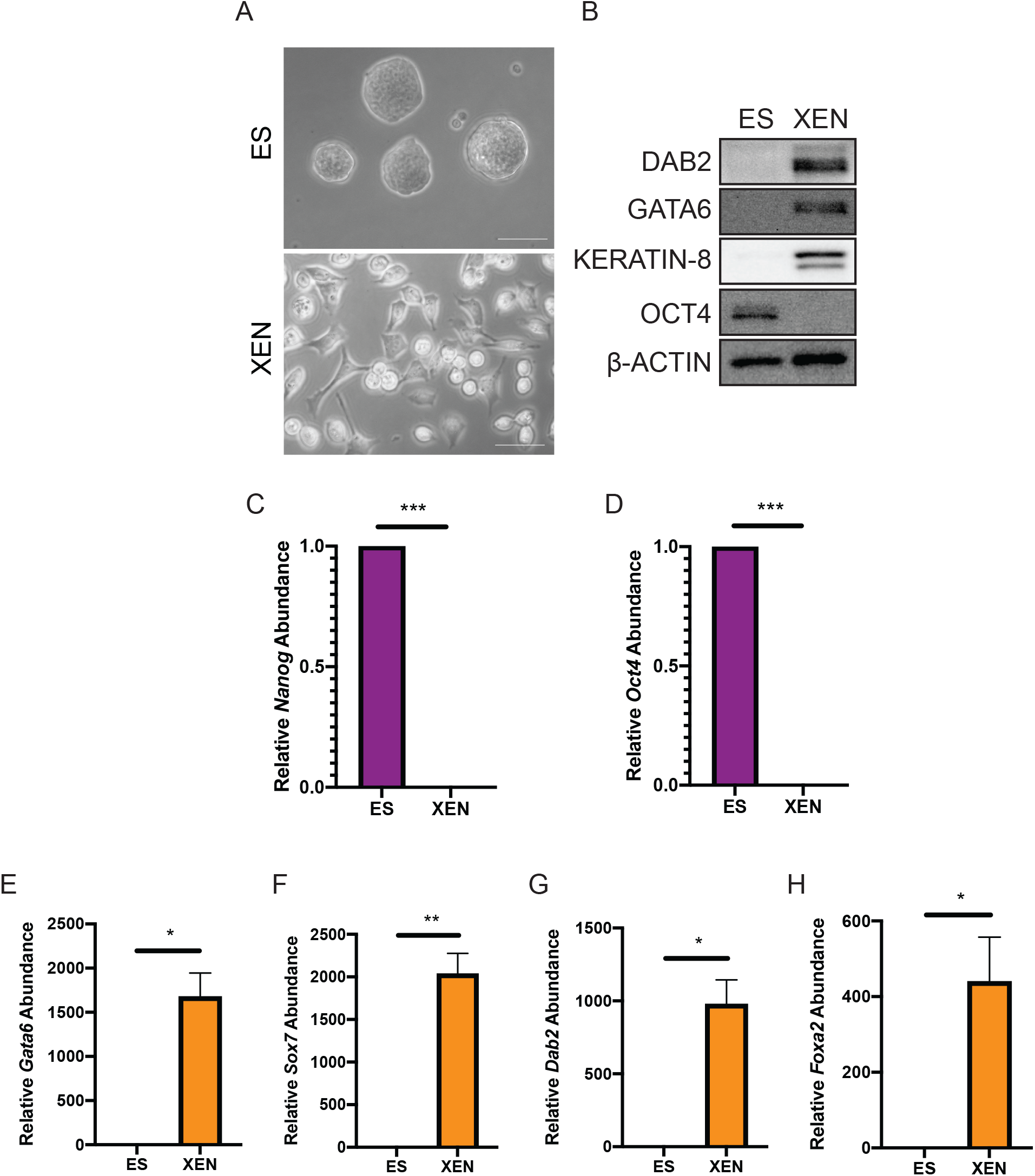
Morphological and molecular analysis of ES and XEN cells in vitro. (A) A representative phase contrast micrograph of ES and XEN cells cultured under maintenance conditions under free-feeder conditions. (B) Immunoblot analysis of OCT4 (pluripotency marker) and DAB2, GATA6, and KERATIN-8 (extraembryonic endoderm markers). β-ACTIN served as a loading control. (C-D) Bar graph of expression level of *Oct4* and *Nanog* in ES and XEN cells under maintenance conditions. *Rpl14* was used a constitutive gene for qRT-PCR. (n = three biological replicates, \*\*\**P* <0.001). (E-H) Bar graph of expression level of *Gata6*, *Sox7*, *Dab2*, and *Foxa2* in ES and XEN cells under maintenance conditions. *Rpl14* was used a constitutive gene for qRT-PCR. (n = three biological replicates, \**P* <0.05, \*\**P* <0.01).

**Figure S2.**
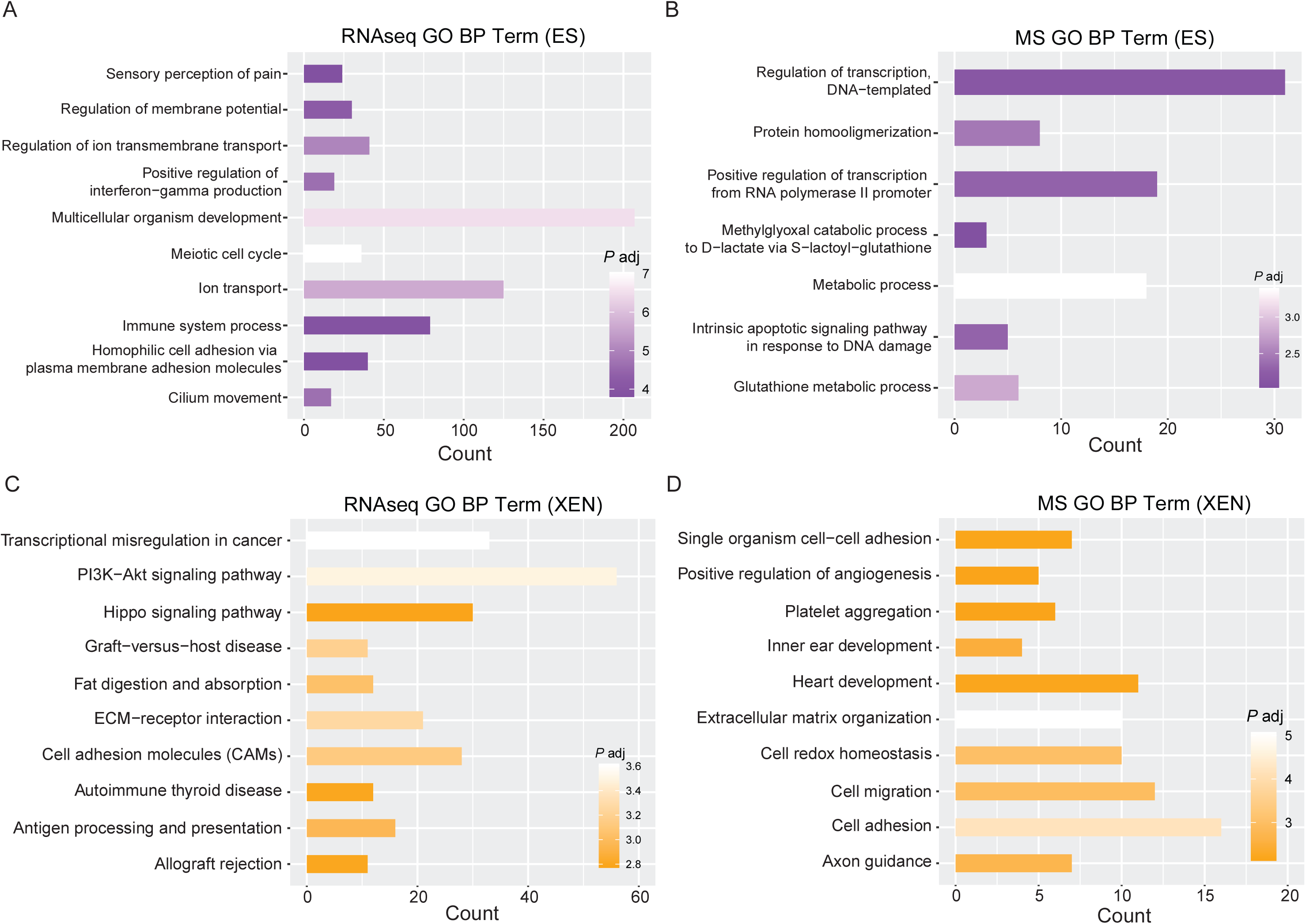
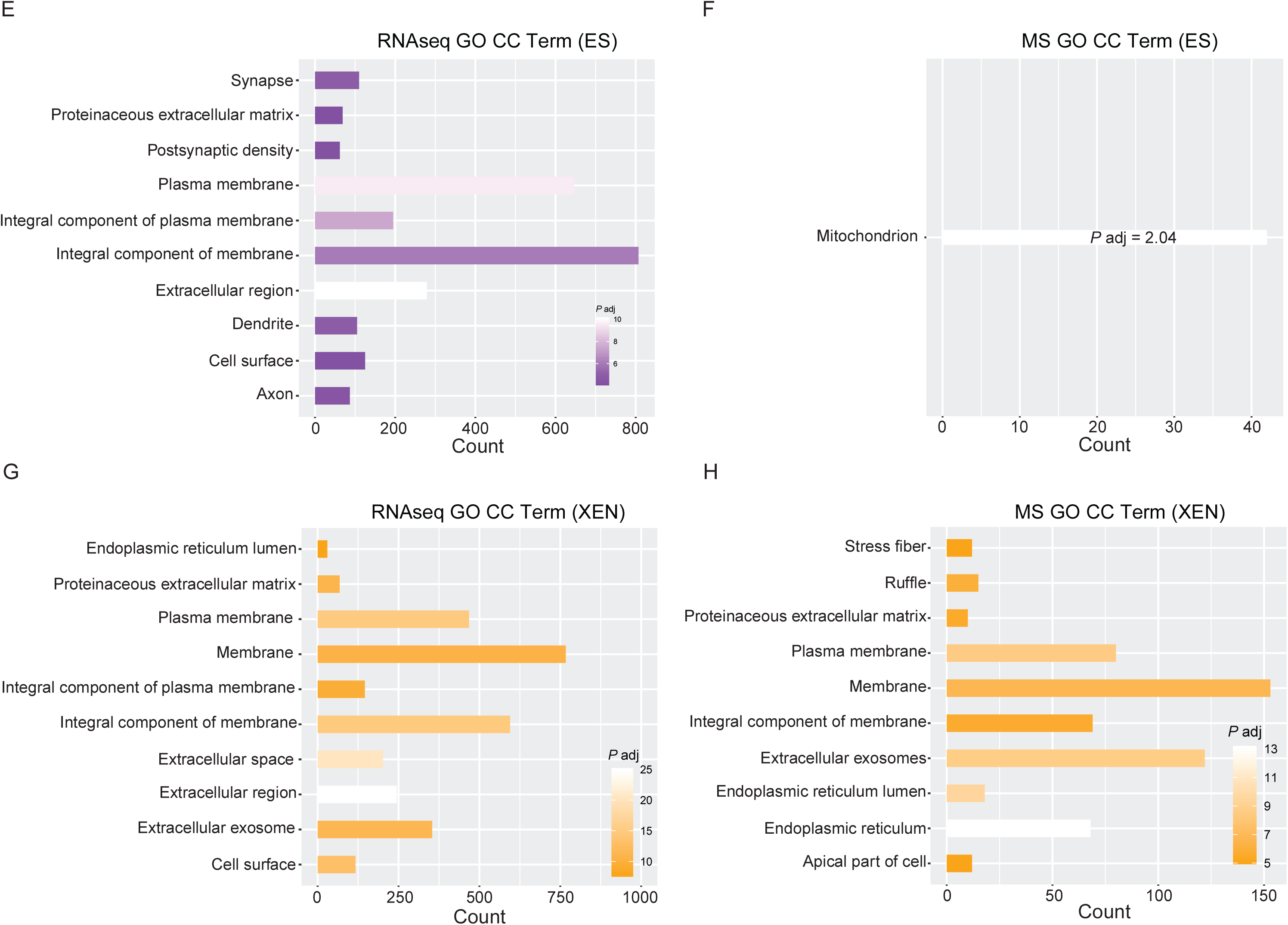
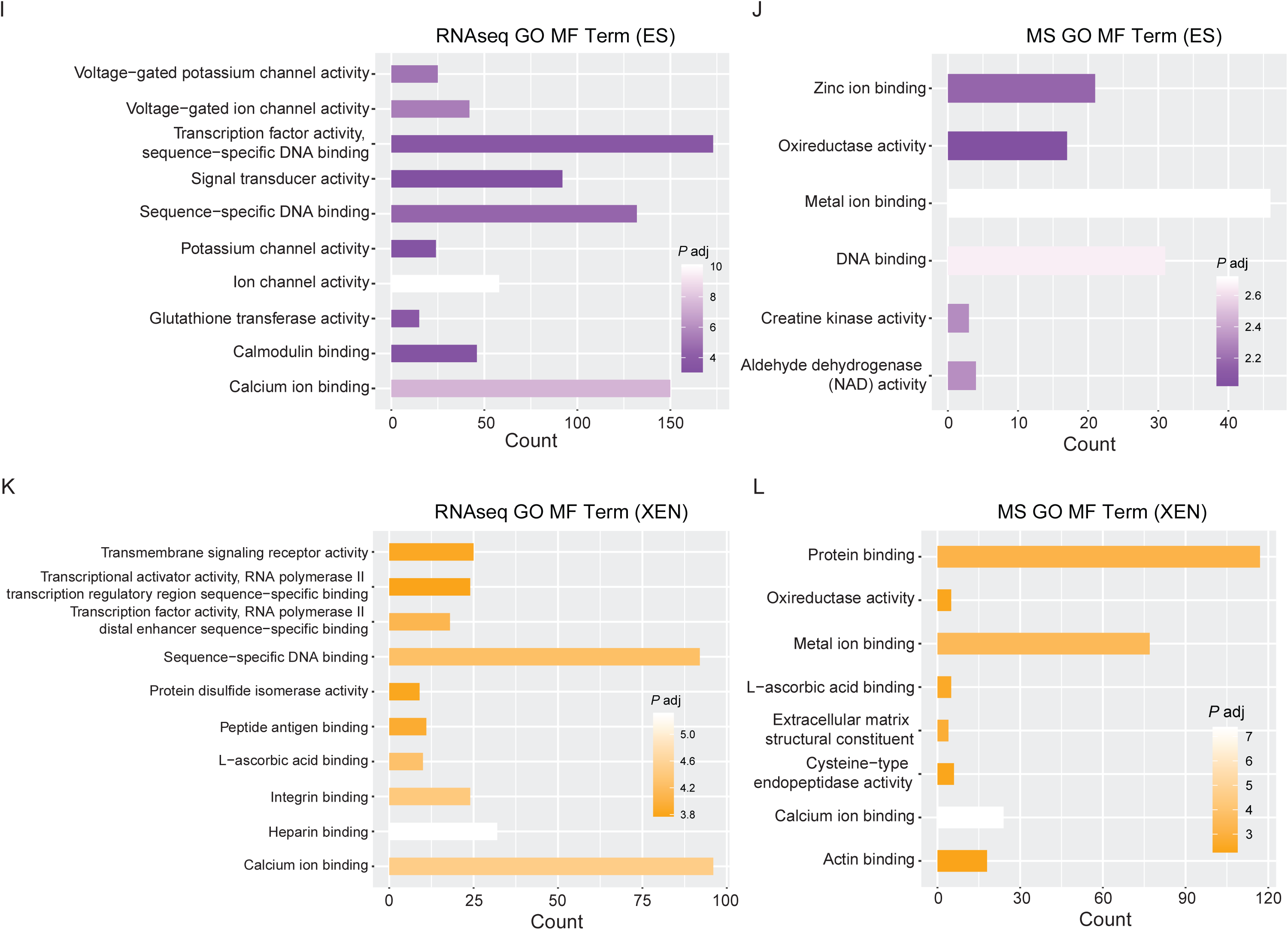
GO term analysis of ES and XEN cells. (A-B) Bar graph representing top 10 enriched GO biological processes terms in ES cells from transcriptomic and proteomic dataset. (C-D) Bar graph representing top 10 enriched GO biological processes terms in XEN cells from transcriptomic and proteomic dataset. (E-F) Bar graph representing top 10 enriched GO cellular compartments terms in ES cells from transcriptomic and proteomic dataset. (G-H) Bar graph representing top 10 enriched GO cellular compartments terms in XEN cells from transcriptomic and proteomic dataset. (I-J) Bar graph representing top 10 enriched GO molecular functions terms in ES cells from transcriptomic and proteomic dataset. (K-L) Bar graph representing top 10 enriched GO molecular functions terms in XEN cells from transcriptomic and proteomic dataset.

**Figure S3.**
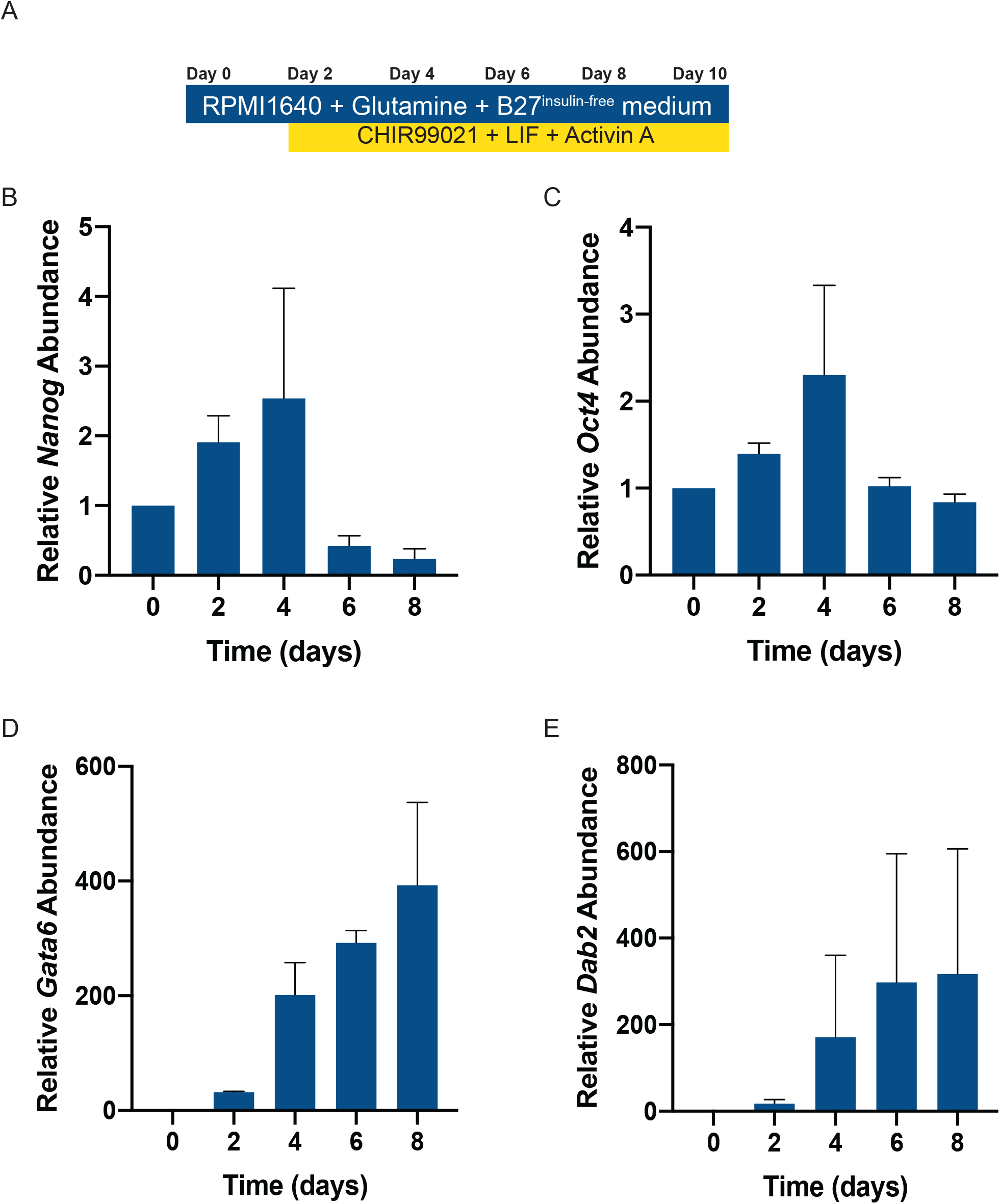
In vitro induction of ES cells towards the XEN lineage. (A) Schematic diagram of the methodology of cXEN induction *in* vitro. (B-C) Bar graph of expression level of *Nanog* and *Oct4* in ES cells induced towards the XEN lineage over 8 days. *Rpl14* was used a constitutive gene for qRT-PCR. (n = three biological replicates). (D-E) Bar graph of expression level of *Gata6* and *Dab2* in ES cells induced towards the XEN lineage over 8 days. *Rpl14* was used a constitutive gene for qRT-PCR. (n = three biological replicates).

## Notes

https://www.ncbi.nlm.nih.gov/geo/query/acc.cgi?acc=GSE159855

### Competing Interest Statement

The authors have declared no competing interest.

### Summary of Updates

Changes were made to update author affiliations only. No other changes were made.

## References

Anderson, K.G.V., Hamilton, W.B., Roske, F.V., Azad, A., Knudsen, T.E., Canham, M.A., Forrester, L.M., and Brickman, J.M. (2017). Insulin fine-tunes self-renewal pathways governing naive pluripotency and extra-embryonic endoderm. Nat Cell Biol 19, 1164–1177.

Baksh, S.C., Todorova, P.K., Gur-Cohen, S., Hurwitz, B., Ge, Y., Novak, J.S.S., Tierney, M.T., Dela Cruz-Racelis, J., Fuchs, E., and Finley, L.W.S. (2020). Extracellular serine controls epidermal stem cell fate and tumour initiation. Nat Cell Biol 22, 779–790.

Brons, I.G., Smithers, L.E., Trotter, M.W., Rugg-Gunn, P., Sun, B., Chuva de Sousa Lopes, S.M., Howlett, S.K., Clarkson, A., Ahrlund-Richter, L., Pedersen, R.A., et al. (2007). Derivation of pluripotent epiblast stem cells from mammalian embryos. Nature 448, 191–195.

Cho, L.T., Wamaitha, S.E., Tsai, I.J., Artus, J., Sherwood, R.I., Pedersen, R.A., Hadjantonakis, A.K., and Niakan, K.K. (2012). Conversion from mouse embryonic to extra-embryonic endoderm stem cells reveals distinct differentiation capacities of pluripotent stem cell states. Development 139, 2866–2877.

Choi, J., Seo, B.J., La, H., Yoon, S.H., Hong, Y.J., Lee, J.H., Chung, H.M., Hong, K., and Do, J.T. (2020). Comparative analysis of the mitochondrial morphology, energy metabolism, and gene expression signatures in three types of blastocyst-derived stem cells. Redox Biol 30, 101437.

Comes, S., Gagliardi, M., Laprano, N., Fico, A., Cimmino, A., Palamidessi, A., De Cesare, D., De Falco, S., Angelini, C., Scita, G., et al. (2013). L-Proline induces a mesenchymal-like invasive program in embryonic stem cells by remodeling H3K9 and H3K36 methylation. Stem Cell Reports 1, 307–321.

Cooper, T.T., Sherman, S.E., Kuljanin, M., Bell, G.I., Lajoie, G.A., and Hess, D.A. (2018). Inhibition of Aldehyde Dehydrogenase-Activity Expands Multipotent Myeloid Progenitor Cells with Vascular Regenerative Function. Stem Cells 36, 723–736.

Davidson, K.C., Mason, E.A., and Pera, M.F. (2015). The pluripotent state in mouse and human. Development 142, 3090–3099.

De Saedeleer, C.J., Copetti, T., Porporato, P.E., Verrax, J., Feron, O., and Sonveaux, P. (2012). Lactate activates HIF-1 in oxidative but not in Warburg-phenotype human tumor cells. PLoS One 7, e46571.

Dobrovolny, P.L., and Bess, D. (2011). Optimized PCR-based detection of mycoplasma. J Vis Exp.

Evans, M.J., and Kaufman, M.H. (1981). Establishment in culture of pluripotential cells from mouse embryos. Nature 292, 154–156.

Fischer, B., and Bavister, B.D. (1993). Oxygen tension in the oviduct and uterus of rhesus monkeys, hamsters and rabbits. J Reprod Fertil 99, 673–679.

Folmes, C.D., Nelson, T.J., Martinez-Fernandez, A., Arrell, D.K., Lindor, J.Z., Dzeja, P.P., Ikeda, Y., Perez-Terzic, C., and Terzic, A. (2011). Somatic oxidative bioenergetics transitions into pluripotency-dependent glycolysis to facilitate nuclear reprogramming. Cell Metab 14, 264–271.

Fujikura, J., Yamato, E., Yonemura, S., Hosoda, K., Masui, S., Nakao, K., Miyazaki Ji, J., and Niwa, H. (2002). Differentiation of embryonic stem cells is induced by GATA factors. Genes Dev 16, 784–789.

Gatie, M.I., and Kelly, G.M. (2018). Metabolic profile and differentiation potential of extraembryonic endoderm-like cells. Cell Death Discov 4, 42.

Gu, W., Gaeta, X., Sahakyan, A., Chan, A.B., Hong, C.S., Kim, R., Braas, D., Plath, K., Lowry, W.E., and Christofk, H.R. (2016). Glycolytic Metabolism Plays a Functional Role in Regulating Human Pluripotent Stem Cell State. Cell Stem Cell 19, 476–490.

Halestrap, A.P. (1975). The mitochondrial pyruvate carrier. Kinetics and specificity for substrates and inhibitors. Biochem J 148, 85–96.

He, X., Chi, G., Li, M., Xu, J., Zhang, L., Song, Y., Wang, L., and Li, Y. (2020). Characterisation of extraembryonic endoderm-like cells from mouse embryonic fibroblasts induced using chemicals alone. Stem Cell Res Ther 11, 157.

Hogan, B.L., Cooper, A.R., and Kurkinen, M. (1980). Incorporation into Reichert's membrane of laminin-like extracellular proteins synthesized by parietal endoderm cells of the mouse embryo. Dev Biol 80, 289–300.

Houghton, F.D., Humpherson, P.G., Hawkhead, J.A., Hall, C.J., and Leese, H.J. (2003). Na+, K+, ATPase activity in the human and bovine preimplantation embryo. Dev Biol 263, 360–366.

Huang da, W., Sherman, B.T., and Lempicki, R.A. (2009a). Bioinformatics enrichment tools: paths toward the comprehensive functional analysis of large gene lists. Nucleic Acids Res 37, 1–13.

Huang da, W., Sherman, B.T., and Lempicki, R.A. (2009b). Systematic and integrative analysis of large gene lists using DAVID bioinformatics resources. Nat Protoc 4, 44–57.

Hui, S., Ghergurovich, J.M., Morscher, R.J., Jang, C., Teng, X., Lu, W., Esparza, L.A., Reya, T., Le, Z., Yanxiang Guo, J., et al. (2017). Glucose feeds the TCA cycle via circulating lactate. Nature 551, 115–118.

Kunath, T., Arnaud, D., Uy, G.D., Okamoto, I., Chureau, C., Yamanaka, Y., Heard, E., Gardner, R.L., Avner, P., and Rossant, J. (2005). Imprinted X-inactivation in extra-embryonic endoderm cell lines from mouse blastocysts. Development 132, 1649–1661.

Langmead, B., and Salzberg, S.L. (2012). Fast gapped-read alignment with Bowtie 2. Nat Methods 9, 357–359.

Latham, T., Mackay, L., Sproul, D., Karim, M., Culley, J., Harrison, D.J., Hayward, L., Langridge-Smith, P., Gilbert, N., and Ramsahoye, B.H. (2012). Lactate, a product of glycolytic metabolism, inhibits histone deacetylase activity and promotes changes in gene expression. Nucleic Acids Res 40, 4794–4803.

Li, B., and Dewey, C.N. (2011). RSEM: accurate transcript quantification from RNA-Seq data with or without a reference genome. BMC Bioinformatics 12, 323.

Love, M.I., Huber, W., and Anders, S. (2014). Moderated estimation of fold change and dispersion for RNA-seq data with DESeq2. Genome Biol 15, 550.

Mali, P., Chou, B.K., Yen, J., Ye, Z., Zou, J., Dowey, S., Brodsky, R.A., Ohm, J.E., Yu, W., Baylin, S.B., et al. (2010). Butyrate greatly enhances derivation of human induced pluripotent stem cells by promoting epigenetic remodeling and the expression of pluripotency-associated genes. Stem Cells 28, 713–720.

Mathieu, J., and Ruohola-Baker, H. (2017). Metabolic remodeling during the loss and acquisition of pluripotency. Development 144, 541–551.

McDonald, A.C., Biechele, S., Rossant, J., and Stanford, W.L. (2014). Sox17-mediated XEN cell conversion identifies dynamic networks controlling cell-fate decisions in embryo-derived stem cells. Cell Rep 9, 780–793.

Mulvey, C.M., Schroter, C., Gatto, L., Dikicioglu, D., Fidaner, I.B., Christoforou, A., Deery, M.J., Cho, L.T., Niakan, K.K., Martinez-Arias, A., et al. (2015). Dynamic Proteomic Profiling of Extra-Embryonic Endoderm Differentiation in Mouse Embryonic Stem Cells. Stem Cells 33, 2712–2725.

Niakan, K.K., Schrode, N., Cho, L.T., and Hadjantonakis, A.K. (2013). Derivation of extraembryonic endoderm stem (XEN) cells from mouse embryos and embryonic stem cells. Nat Protoc 8, 1028–1041.

Nichols, J., Evans, E.P., and Smith, A.G. (1990). Establishment of germ-line-competent embryonic stem (ES) cells using differentiation inhibiting activity. Development 110, 1341–1348.

Nowotschin, S., Hadjantonakis, A.K., and Campbell, K. (2019). The endoderm: a divergent cell lineage with many commonalities. Development 146.

Paca, A., Seguin, C.A., Clements, M., Ryczko, M., Rossant, J., Rodriguez, T.A., and Kunath, T. (2012). BMP signaling induces visceral endoderm differentiation of XEN cells and parietal endoderm. Dev Biol 361, 90–102.

Parenti, A., Halbisen, M.A., Wang, K., Latham, K., and Ralston, A. (2016). OSKM Induce Extraembryonic Endoderm Stem Cells in Parallel to Induced Pluripotent Stem Cells. Stem Cell Reports 6, 447–455.

Polanski, R., Hodgkinson, C.L., Fusi, A., Nonaka, D., Priest, L., Kelly, P., Trapani, F., Bishop, P.W., White, A., Critchlow, S.E., et al. (2014). Activity of the monocarboxylate transporter 1 inhibitor AZD3965 in small cell lung cancer. Clin Cancer Res 20, 926–937.

Ryu, J.M., and Han, H.J. (2011). L-threonine regulates G1/S phase transition of mouse embryonic stem cells via PI3K/Akt, MAPKs, and mTORC pathways. J Biol Chem 286, 23667–23678.

Seo, B.J., Choi, J., La, H., Habib, O., Choi, Y., Hong, K., and Do, J.T. (2020). Role of mitochondrial fission-related genes in mitochondrial morphology and energy metabolism in mouse embryonic stem cells. Redox Biol 36, 101599.

Shimosato, D., Shiki, M., and Niwa, H. (2007). Extra-embryonic endoderm cells derived from ES cells induced by GATA factors acquire the character of XEN cells. BMC Dev Biol 7, 80.

Soprano, D.R., Teets, B.W., and Soprano, K.J. (2007). Role of retinoic acid in the differentiation of embryonal carcinoma and embryonic stem cells. Vitam Horm 75, 69–95.

Sutendra, G., Kinnaird, A., Dromparis, P., Paulin, R., Stenson, T.H., Haromy, A., Hashimoto, K., Zhang, N., Flaim, E., and Michelakis, E.D. (2014). A nuclear pyruvate dehydrogenase complex is important for the generation of acetyl-CoA and histone acetylation. Cell 158, 84–97.

Tobias, I.C., Isaac, R.R., Dierolf, J.G., Khazaee, R., Cumming, R.C., and Betts, D.H. (2018). Metabolic plasticity during transition to naive-like pluripotency in canine embryo-derived stem cells. Stem Cell Res 30, 22–33.

Tsogtbaatar, E., Landin, C., Minter-Dykhouse, K., and Folmes, C.D.L. (2020). Energy Metabolism Regulates Stem Cell Pluripotency. Front Cell Dev Biol 8, 87.

Vacanti, N.M., Divakaruni, A.S., Green, C.R., Parker, S.J., Henry, R.R., Ciaraldi, T.P., Murphy, A.N., and Metallo, C.M. (2014). Regulation of substrate utilization by the mitochondrial pyruvate carrier. Mol Cell 56, 425–435.

Wamaitha, S.E., del Valle, I., Cho, L.T., Wei, Y., Fogarty, N.M., Blakeley, P., Sherwood, R.I., Ji, H., and Niakan, K.K. (2015). Gata6 potently initiates reprograming of pluripotent and differentiated cells to extraembryonic endoderm stem cells. Genes Dev 29, 1239–1255.

Wang, J., Alexander, P., Wu, L., Hammer, R., Cleaver, O., and McKnight, S.L. (2009). Dependence of mouse embryonic stem cells on threonine catabolism. Science 325, 435–439.

Wessel, D., and Flugge, U.I. (1984). A method for the quantitative recovery of protein in dilute solution in the presence of detergents and lipids. Anal Biochem 138, 141–143.

Yanes, O., Clark, J., Wong, D.M., Patti, G.J., Sanchez-Ruiz, A., Benton, H.P., Trauger, S.A., Desponts, C., Ding, S., and Siuzdak, G. (2010). Metabolic oxidation regulates embryonic stem cell differentiation. Nat Chem Biol 6, 411–417.

Ying, Q.L., Wray, J., Nichols, J., Batlle-Morera, L., Doble, B., Woodgett, J., Cohen, P., and Smith, A. (2008). The ground state of embryonic stem cell self-renewal. Nature 453, 519–523.

Yuan, M., Breitkopf, S.B., Yang, X., and Asara, J.M. (2012). A positive/negative ion-switching, targeted mass spectrometry-based metabolomics platform for bodily fluids, cells, and fresh and fixed tissue. Nat Protoc 7, 872–881.

Zhang, J., Nuebel, E., Daley, G.Q., Koehler, C.M., and Teitell, M.A. (2012). Metabolic regulation in pluripotent stem cells during reprogramming and self-renewal. Cell Stem Cell 11, 589–595.

Zhou, W., Choi, M., Margineantu, D., Margaretha, L., Hesson, J., Cavanaugh, C., Blau, C.A., Horwitz, M.S., Hockenbery, D., Ware, C., et al. (2012). HIF1alpha induced switch from bivalent to exclusively glycolytic metabolism during ESC-to-EpiSC/hESC transition. EMBO J 31, 2103–2116.

